# Visualisation of translating ribosomes reveals the earliest steps of protein misfolding in human disease

**DOI:** 10.1101/2025.04.25.650147

**Authors:** Elena Plessa, Lien P. Chu, Charity R. Hornby, Julian O. Streit, Alkistis N. Mitropoulou, Sammy H.S. Chan, James A. Irving, David A. Lomas, John Christodoulou, Lisa D. Cabrita

## Abstract

The majority of cellular proteins must adopt a particular three-dimensional structure for function^1^. However, protein folding is a perilous journey due to competing polypeptide misfolding events which result in inactive structures. In this study, we examine the earliest steps of protein misfolding during the biosynthesis of alpha-1-antitrypsin, a secreted plasma protein whose misfolding results in organ disease. Using human cells, we find that, like co-translational protein folding, misfolding, assembly and biosynthesis are interconnected processes. At the molecular level misfolding of alpha-1-antitrypsin is initiated by a molten globule-like folding intermediate formed cotranslationally on the ribosome. The ribosomal complexes subsequently form assemblies by recruiting released proteins, inducing translational arrest. Our data also reveal that a pharmacological chaperone modulates this process. The existence of co- and post-translational (mis)folding and assembly pathways reveals how some proteins form functional complexes, has implications for the pathogenesis of conformational diseases, and suggests novel therapeutic avenues.

---

Correct polypeptide folding is essential for producing biologically active proteins at the concentrations needed to support essential cell functions. However, *in vitro* studies have long-indicated that protein folding can deviate off-pathway towards misfolding and mis-assembly, generating functionally inactive alternative structures and even amorphous aggregates. Within cells, protein folding begins co-translationally on ribosomes^1^. However folding is an imperfect process in which 15-30% of all newly-synthesised proteins are ubiquitinated and degraded^2^. Although the cell has evolved quality control measures to mitigate the hazardous effects of protein misfolding, under extreme conditions, these mechanisms can fail, resulting in cell toxicity and disease.

Indeed, there is an increasing range of conformational diseases^3^, spanning neurodegenerative diseases (e.g., Huntington’s, Alzheimer’s, and Parkinson’s diseases), metabolic and organ diseases which are caused by misfolding-prone proteins which form oligomers, highly-ordered amyloids, and aggregates. Within this grouping is the serpin alpha-1-antitrypsin (AAT), which is one of the most abundant proteins in blood, serving diverse functional roles, with a primary role in protecting the lungs from the proteolytic activity of neutrophil elastase during inflammation^4^. Alpha-1-antitrypsin deficiency is caused by the accumulation of toxic misfolded AAT polymers at the site of synthesis in the endoplasmic reticulum of hepatocytes^5^. These polymers are responsible for the development of liver cirrhosis, fibrosis, and hepatocellular carcinoma to homozygotes, as well as early onset of emphysema due to the depletion of the functional AAT in circulation^6^. AAT is therefore an archetypal model for both protein misfolding and mis-assembly. It is well-established that the propensity of AAT to misfold results from point mutations that divert the protein away from its functional folding pathway. The Z mutation, E342K, is the most severe pathological variant of AAT. This point mutation causes a kinetic folding defect which traps the protein within a misfolding-prone, intermediate structure which subsequently self-associates to form highly-ordered, toxic polymers^7^. As shown *in vitro* using ribosome-nascent chain complexes (RNCs), a strategy which captures protein biosynthesis and folding as “biosynthetic snapshots”, Z-alpha-1-antitrypsin (Z-AAT) folding intermediates begin forming co-translationally on ribosomes, thus impacting how the polypeptide completes its native fold following its release^8^. However, it is unclear how the distinct conformation of Z-AAT NCs on the ribosome compared to the wild-type variant, impact the misfolding and aggregation of the protein observed post-translationally.

Understanding the causative relationship that connects defective protein biosynthesis and misfolding with disease is in its infancy. There are no molecular-level descriptions of how and when proteins begin misfolding co-translationally nor of the pathways and resultant changes in protein structure that result. To explore these open questions, we examined Z-AAT as a relevant model of protein misfolding and disease within cells. We developed a multi-scalar toolkit to provide a molecular basis for analysis that reveals a new mechanism of protein misfolding which begins cotranslationally on stalled ribosomes. As modelled by experimentally reweighted molecular dynamics simulations, a non-native conformational ensemble is formed by ribosome-bound nascent polypeptides. This conformation underpins a novel mode of recruitment of released protein intermediates, which can drive ribosome-bound mis-assemblies. Additionally, we find that energetics intrinsically encoded within the amino acid sequence may contribute in modulating translational arrest of polypeptides. Our findings provide new mechanistic insight into the molecular basis connecting ribosome dysfunction with co-translational protein misfolding and mis-assembly and the role of this relationship in the pathogenesis of conformational diseases.

## Detection of co-translational misfolding and mis-assembled complexes on ribosomes in human cells

Our *in vitro* analysis of AAT biosynthesis^8^ revealed natural pausing of ribosomes which facilitated the co-translational folding of this protein. We used this finding as a basis to initiate studies of the pathological variant, Z-AAT, to test whether co-translational misfolding and mis-assembly on ribosomes could be detected in human cells (Fig.1a). Therefore, we developed approaches that detect ribosome dysfunction and kinetically-trapped intermediates and polymers of Z-AAT that emerge co-translationally. This approach assesses the presence of hallmark features of protein misfolding at the earliest possible time points when aberrant folding might occur. To this end, we created a set of DNA constructs to study the biosynthesis of full-length Z-AAT (i.e., mature form with a stop codon) (Fig.1b) in the ER. We also obtained a “final biosynthetic snapshot^8,9^” of Z-AAT misfolding by fusing an arrested-enhanced variant of Xbp1u^10^ at the C-terminus to generate ribosome-nascent chain complexes (RNC); these engineered RNCs (Fig.1b,c) were used to mimic translational arrest. We developed a doxycycline-inducible method (Extended Data Fig.1) using synchronised human (HEK293 and HeLa) cells to selectively synthesise Z-AAT in the endoplasmic reticulum (or the cytosol^8^). A 5-hour induction period was used to generate a “cellular snapshot” of early biosynthesis within one cell cycle (18 to 24 h), while longer time periods (up to 48 hours) allowed evaluation of protein misfolding during prolonged Z-AAT over-expression.

**Figure 1:**
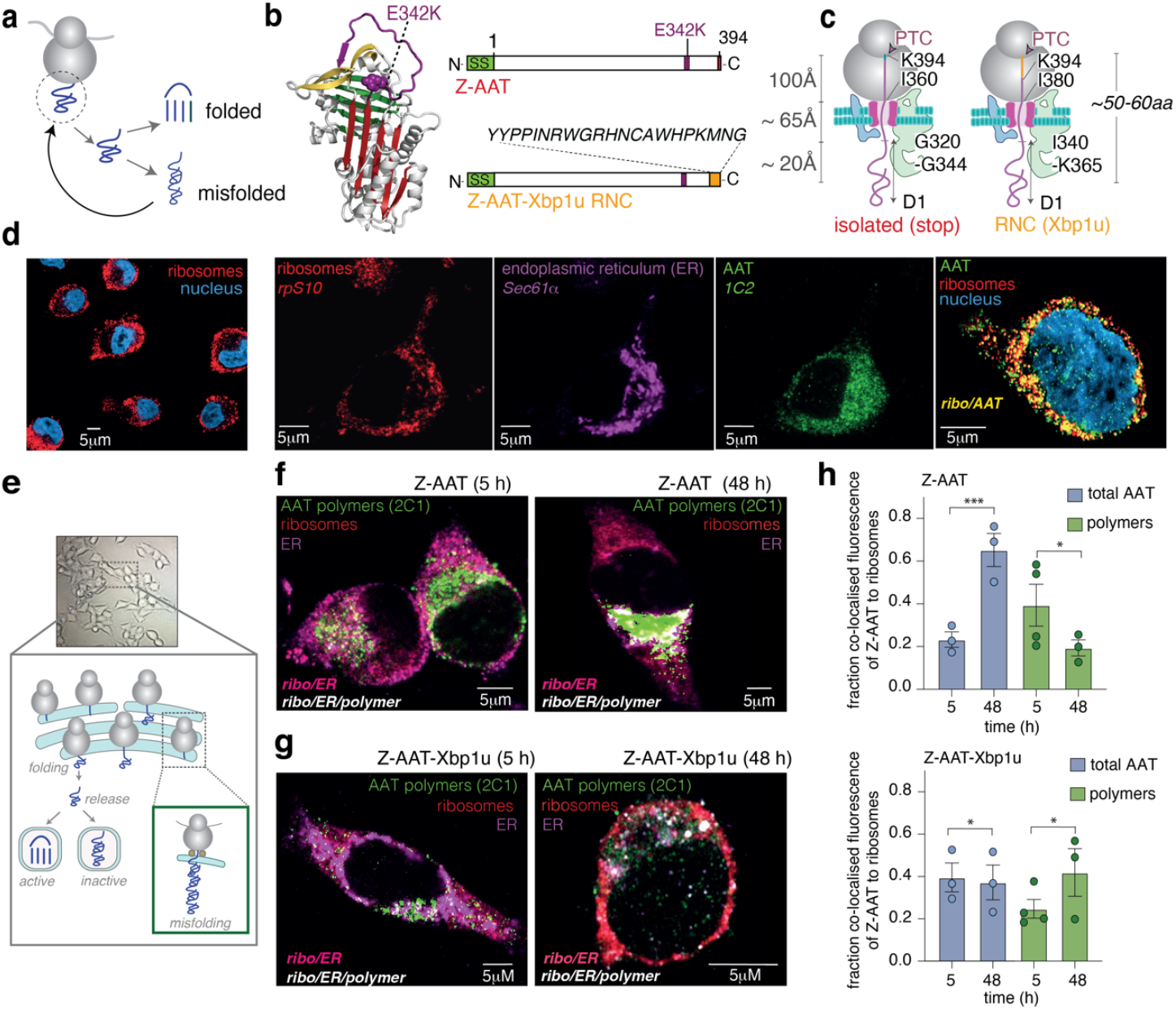
Co-translational misfolding and mis-assembly on translating ribosome in human cells. **(a)** Tracing back the earliest origins of protein misfolding to the ribosome. **(b)** *(left)* Structure of AAT (1QLP) with the Z mutation (E342K) indicated. *(right)* AAT constructs used in this study. **(c)** Schematic of ER-bound RNCs with AAT’s amino acid sequence mapped (predicted). **(d)** Confocal fluorescence microscopy images showing detection of ribosomes (rpS10), Z-AAT (1C2), ER (Sec61α) and the nucleus (DAPI staining). **(e)** Models of AAT folding and misfolding in the ER. **(f)** Confocal microscopy images of isolated Z-AAT polymer formation (2C1) at 5 and 48 hours. Images represent merged ER (Sec61α), ribosome (rpS10) and AAT polymer channels (2C1). **(g)** As described in (f), but showing Xbp1u-arrested Z-AAT RNCs. **(h)** Quantification of co-localisation of total Z-AAT (1C2) and Z-AAT polymers (2C1) with 80S ribosomes (M2, AAT channel) at 5 and 48 hours (SEM, n = 4, measurements taken across different transfections).

We initially addressed whether we could visualise the biosynthesis and polymerisation of Z-AAT via multiplex immunofluorescence (Figs.1d,f, Extended Data Figs.2a-d) in fixed cells using antibodies recognising the ER (Sec61α), 80S ribosomes (eS10), and either total Z-AAT (monomers and polymers, 1C2 antibody) or its polymeric form (2C1 antibody^11^) (Fig.1d,f, Extended Data Fig. 2c). At 5 hours, Z-AAT was broadly distributed throughout the ER with visible colocalisation to ER-bound ribosomes (34 ± 5.4%), as validated by a Manders correlation coefficient (MCC) analysis (Figs. 1h, Extended Data Fig. 3a). The MCC analysis validated the co-localisation observed between ribosomes and the ER during the expression of Z-AAT over time (representative image, Fig.1f, pink puncta). At 5 hours, the majority of the fluorescence signal from the ER overlapped with ribosome-specific fluorescence (79 ± 3 %, Extended Data Fig.3a), as anticipated given the proximity between the ER and ribosomes. Following 48 hours of prolonged expression, fewer ER-bound ribosomes (26 ± 3 %, Extended Data Fig.3a) were observed, suggesting potential ER stress^12^.

Next, we probed for the presence of Z-AAT polymers: there were clear and discernible ribosome-bound polymers in the ER (Fig.1f, white puncta) at 5 hours. We note these images were collected with the ER present within the focal plane, which enables the ribosome-bound polymers to be observed at a resolution of ∼200 nm in the x-y plane (Figs.1d,f,g). By 48 hours, there was a visible decrease in polymers indicating their release from the ribosome (MCC values decreased from 39± 4.8% to 28± 4.6%, Fig.1h & Extended Data Fig.3b)). These latter results also coincided with a substantial population of Z-AAT retained in the ER (26± 7.3%, Fig.1f & Extended Data Figs. 2c,3b) and on ER-bound ribosomes (50 ± 8.2%, Extended Data Fig.3b). Similar results were obtained from the Xbp1u-arrested Z-AAT, which assesses mis-assembly (i.e., AAT polymerisation) on RNCs that were specifically designed to stall (Fig.1g white puncta); for this construct, Z-AAT expression remained constant (Extended Data Fig.2d), and generally, more ribosome-bound polymers were observed over time (Figs.1g,h). Overall, the fluorescence images provide *in vivo* evidence that protein misfolding and mis-assembly initiates co-translationally on arrested ribosomes.

**Figure 2:**
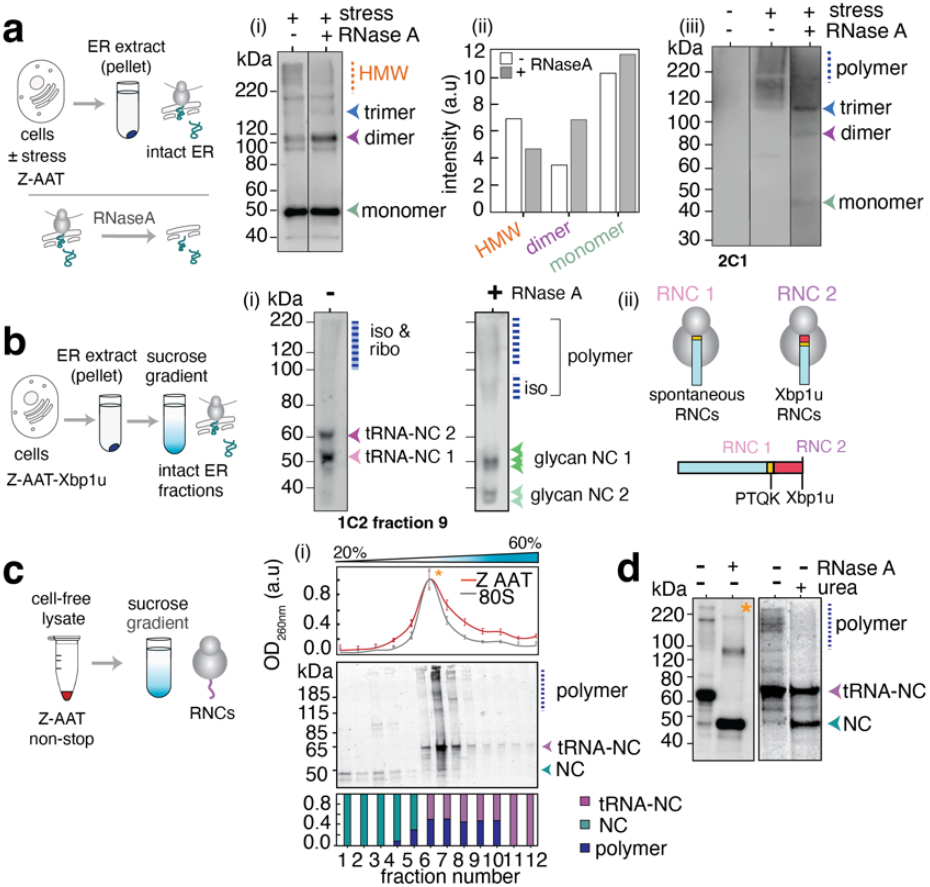
*In vivo* and *in vitro* biochemical characterisation of co-translational misfolding, and mis-assembly formation on ribosomes. **(a)** *(upper)* Schematic image showing purification of ER-bound Z-AAT RNCs from HEK293 cells. *(lower)* RNaseA treatment releases ribosome-bound polymers and dimers located inside the ER of HEK293 cells. (i) Naturally-paused ER-bound ribosomes expressing Z-AAT (1C2 detection) recovered from intact ER (± RNase A). Cells were subjected to ER stress for 48 hours (n=2). (ii) Bar plot showing the species identified from gel (i) (±RNase A). (iii) Ribosome-bound polymers of Z-AAT (2C1 detection) following 48 hours of expression with and without ER stress, ± RNase A (n=2). **(b)** Schematic image showing the purification of ER-bound Z-AAT-Xbp1u RNCs from HEK293 cells. (i) Sucrose gradient purified ER-bound, Z-AAT-Xbp1u RNCs (fraction 9, ±RNase A) (n=4). (ii) Schematic image of RNC species formed during cell stress derived from analysis of Xbp1u-induced arrest (see text). **(c)** Schematic image showing the purification of (non-stop) Z-AAT RNCs expressed in RRL (cell-free). (i) *(upper)* Overlay of sucrose gradient profiles of Z-AAT RNCs and (empty) 80S ribosomes (UV detection). *(middle)* Sucrose gradient fractionation of Z-AAT RNCs monitored by ^35^S-Met autoradiography on semi-denaturing PAGE (n=3) (*lower*) Densitometry analysis of fractions (n=3). **(d)** Z-AAT RNCs following a 22-hour incubation in buffer ±RNase A and 8M urea treatment. All gels are representative of biological repeats.

## AAT forms ribosome-bound polymers as a result of translational arrest

To study a potential correlation between translational arrest and protein misfolding, we examined AAT biosynthesis in the ER within HEK293 cells (Fig.2a & Extended Data Fig. 4a) by applying a synchronised approach exploiting the growth of serum-starved cells and harringtonine-treated ribosomes (Extended Data Fig.1a). An analysis of ER extracts taken from a small population of these cells (10^5^) revealed a fraction of glycosylated Z-AAT retained after 3 hours (Extended Data Fig. 4a). These data were consistent with our initial confocal analysis (Extended Data Fig.2) and indicate that AAT successfully translocates into the ER where folding and glycosylation occurs, and thus demonstrating that we can generate “cellular snapshots” of early biosynthesis and dissect this in detail. The same samples also revealed a low population (∼3% of total AAT) of RNCs at 24 hours (Extended Data Fig.4a(i)) with a molecular weight (∼120kDa) consistent with a ribosome-bound Z-AAT dimer, which downshifts after RNaseA treatment (Extended Data Fig.4a (iii), *see later*). Overall, these findings indicate that misfolded intermediates arise co-translationally and remain localised to the ER where they can participate in dimerisation events on the ribosome.

Next, we tested the impact of chemically-induced ER stress on Z-AAT expression (Fig.2a), as a way of enhancing the observation of co-translational misfolding within a compromised cellular environment. We found a modest increase in dimer formation at 9 hours (Extended Data Fig.4a(ii)) under these conditions. Since this result was indicative of enhanced formation Z-AAT polymers forming on ribosomes, we followed up on this observation in depth. Following 48-hour expression of Z-AAT in a larger cell population (∼10^7^) under stressed and non-stressed conditions, the ER showed a substantial accumulation of high molecular weight (HMW) multimers (>220kDa, Fig.2a(i)). Of the observable multimers, ∼2% were found to be ribosome-bound as evident from the downshift of the HMW species into dimers which is the minimum unit of polymer formation in these stressed cells (Fig.2a & Extended Data Fig.4b). These HMW species (formed during both stressed and unstressed conditions) were detected by the 2C1 AAT polymer antibody (Fig.2a(iii)).

Taken as an average at this 48-hour cellular snapshot, the presence of ER-resident, ribosome-bound dimers reveal processes involving translational stalling and co-translational misfolding and which are presumably initiated by retention of Z-AAT protein in the ER.

To understand the origin of the spontaneous and misfolding-prone RNCs, we studied the expression of Xbp1u-stalled Z-AAT RNCs where C-terminal Xbp1u motif forces ribosomes to stall. These designed RNCs offered a way to independently assess stalling and mis-assembly. We used a sucrose gradient to purify the ER extract and examined the resulting ER-bound RNCs (Fig. 2b). After 30 mins of expression, two distinct RNC sizes were observed (Fig.2b, & Extended Data Figs.4c,d). These sizes were consistent with stalling at the stop codon and the Xbp1u arrest motif, respectively (Fig.2b & Extended Data Fig.4d). Compared to Z-AAT (with a stop codon), however, there were fewer released nascent chains (NCs) observed after 24h in Xbp1u-arrested AAT. In addition, there were no ribosome-bound dimers, suggesting that NC release and ribosome-bound polymer formation are correlated. Additionally, extensive protein heterogeneity was observed including glycosylated species, full-length released Z-AAT NCs, truncated fragments (*ca*. 40kDa, Fig.2d), and ribosome-bound Z-AAT (likely at least partially glycosylated, Extended Data Fig.4f, and *see later*). There was a mix of released and ribosome-bound polymers (Figs.2b, & Extended Data Figs.4d,e), which is reflective of the dynamic cellular environment captured within this snapshot (Fig.2e). As demonstrated with the Z-Xbp1u-RNCs, the Xbp1u motif offers a cell-based system that reports on transient translational pausing events (Fig.2b) (e.g., that occur at stop codons^13^, including for AAT *in vitro*^8^). This effect likely results from local stress induced by forced ribosome arrest due to the Xbp1u stalling sequence, which enhances the frequency of otherwise naturally-occurring translational pausing caused by cell stress.

Overall, the capacity to extract intact ER harbouring bound ribosomes shows that translational arrest and protein misfolding are intimately linked processes. These processes can be exacerbated by misfolding-prone NC during translation; left uncontrolled, this can orchestrate cotranslational (mis)assembly formation on translating ribosomes, as observed here with Z-AAT.

### Ribosome-bound polymers as observed during real-time biosynthesis *in vitro*

To formulate a molecular-based understanding of how ribosome-bound polymers form, we studied the biosynthesis of Z-AAT in ER-like microsomes using *in vitro* coupled transcription-translation in rabbit reticulocyte lysate (RRL) and ^35^S methionine autoradiography (Fig.2c & Extended Data Figs. 5c,d). A range of species were observed over time including full-length glycosylated Z-AAT RNCs (65kDa) (Extended Data Fig.5d), released NCs (50 kDa), and a high molecular weight (HMW) banding pattern consistent with polymers^8^ (Extended Data Figs. 5c,d).

To study the link between ribosome pausing/arrest and polymerisation, we quantified the population of AAT species (RNCs, NCs, polymers) that formed over 60 minutes (Extended Data Fig.5f) by applying an ATCA-synchronised approach (coupled translation, no microsomes)^8^. The formation of full-length Z-AAT RNCs peaked at 15 min alongside the increased appearance of released NCs (full-length 47kDa and truncated, 30 kDa) and polymers (after 13 min) (Extended Data Fig.5f). A comparison between Z-AAT and the wild-type AAT revealed that the Z-AAT RNC population peaks 5 min earlier relative to the wild-type (15 min versus 20 min) (Extended Data Fig.5f). This slight difference in transient pausing between the wild-type AAT and Z-AAT may potentially reflect how quickly (ribosome-bound) polymers form.

Next, we characterised the Z polymer species formed during cell-free biosynthesis (no microsomes) using sucrose-density gradients. This analysis showed that the predominant species were full-length NCs and polymers on monosomes, while ribosome-bound dimers were also observed (Fig.2c & Extended Data Fig.5g). When the same reactions were undertaken via uncoupled translation (with exogenous mRNA), polysomes harbouring NCs of different lengths were evident (Extended Data Fig.5b and discussed later) and so too were ribosome-bound polymers. Since equivalent results were also obtained when using either stop or non-stop DNA constructs (Fig. 2c & Extended Data Fig.5g), we continued characterisation with the non-stop construct (and coupled translation) to enhance the population of ribosome-bound polymers.

We used a urea treatment to unfold the ribosome-bound polymers, which resulted in two distinct populations, ribosome-bound monomers and released Z-AAT NCs (Fig.2d), indicating a recruitment process where released NCs can be assembled onto translating ribosomes. This recruitment was explored further by incubating Z-AAT RNCs in aqueous buffer (25°C, 24 hours, Extended Data Fig.6a) to monitor NCs stochastically releasing from the ribosomal complexes. We detected such NC release occurring as early as 5 hours (by Western blot), and by 22 hours released NCs accounted for ∼30 % of observable species. Meanwhile, there was a concomitant increase in ribosome-bound polymers (Extended Data Fig.6a).

### Direct observation of ribosome-bound AAT polymers using electron microscopy

To establish a structural understanding of the ribosome-bound polymers, we used negative stain transmission electron microscopy. First, we visualised Z-AAT RNCs formed following 20 min of coupled synthesis in RRL (stop codon DNA construct, Fig.3a & Extended Data Fig.6b-f). The images were of excellent quality showing a clear uranyl-acetate staining (i.e., dark border) of 80S ribosomes in both monosomes and polysomes (Fig.3a). Protruding from many monosomes were chains of different lengths reminiscent of AAT polymers (Fig.3a). We also observed released polymers (42-400nm) (Fig.3a & Extended Data Fig.6e). The non-stop Z-AAT RNCs (Extended Data Fig.6c) showed equivalent results by negative-stain EM, and their generally higher population enabled features to be more definitively resolved (Fig.3b), including a clear observation of the large 60S subunit, confirming emergence of the polymer chain from the ribosome exit tunnel (Fig.3b). The ribosome-bound polymers varied in length (25-50 nm), and showed a beads-on-a-string morphology (Fig.3b & Extended Data Fig.6e). The monomer units were 4 ± 1 nm in length, revealing that ribosome-bound polymers were comprised of 6-13 AAT monomers per ribosome (Fig.3b & Extended Data Fig.6b-f). Additionally, the accompanying increase in molecular weight of the RNC (from 47 kDa to 282-611 kDa) does not compromise the overall integrity of the RNC (Fig.3a,b & Extended Data Fig.6c,f). Longer, released polymers of up to 400 nm long (i.e., 100 AAT monomer units) (Fig. 3a & Extended Data Fig.6e,f) were also observed, suggesting that a size limit for co-translational assembly exists.

**Figure 3:**
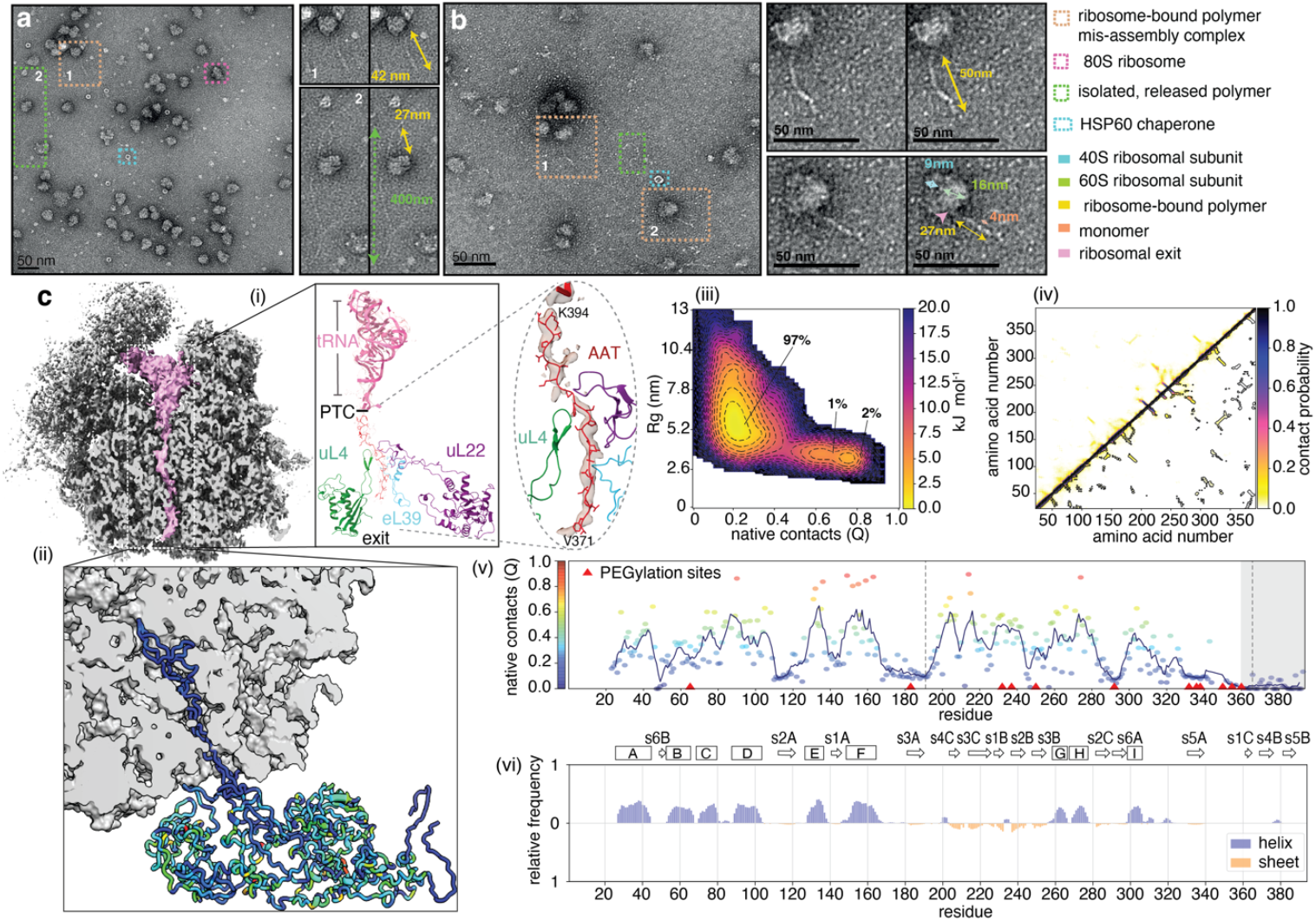
A ribosome-bound co-translational folding intermediate drives co-post assembly mechanism on the ribosome. Micrographs of uranyl acetate-stained Z-AAT RNCs produced in RRL: **(a)** Z-AAT samples following 20 minutes of biosynthesis in RRL. *(inset)* Magnified views of RNC species and polymers. **(b)** Purified (non-stop) Z-AAT RNCs following incubation at room temperature for 22 hours. *(inset)* Magnified views of 2 RNC species. **(c)** (i) A 3.7 Å cryo-EM map cross-section of wild-type AAT RNC (non-stop), with the P-site tRNA and bound AAT NC electron densities highlighted in pink. *(inset)* a magnified view of the ribosomal exit tunnel showing the lysine tRNA (pink) and AAT NC (red) fitted inside the electron density of the RNC map. (ii) Structure of wild-type AAT’s ribosome-bound, co-translational intermediate ensemble derived from experimentally-reweighted MD simulations. Three representative conformations with significant populations (3.4%) are shown and coloured by fraction of native contacts (Q). (iii) A two-dimensional energy landscape depicting the distribution of the radius of gyration (Rg) values versus fraction of native contacts (Q), as observed for the conformational ensemble following reweighting. (iv) A contact map for the wild AAT RNC. No long-range contacts are observed, consistent with an expanded and highly dynamic heterogeneous conformation. (v) A weighted ensemble average (5-residue window size) showing the fraction of native contacts (Q) per residue observed in the wild-type AAT RNC. Red triangles indicate single cysteine positions used to generate protection factors, the dashed lines indicate the 2 proteolytically-resistant fragments^8^, and the shaded region are residues expected to be in the exit tunnel. (vi) A per-residue plot of secondary structure of the wild-type AAT RNC.

The effect of NC length on co-translational polymerisation formation was studied using a set of C-terminal truncations (ranging from 50 to 300 amino acids). All the constructs tested induced translational pausing (Extended Data Fig.6k, and see later), but the minimum length at which any form of HMW species was observed occurred between 200 and 250 amino acids (D1 to A250) (Extended Data Fig.6k). These results suggest that polymer formation, or at least, a form of cotranslational assembly, requires a minimum NC length for structure to form; for AAT at least, this is 200 amino acids (Extended Data Fig.6k). This length coincides with the formation of a compacted N-terminal domain (see Ref.^8^ and see later). Coincidentally, although disease-causing polymers form via a full-length AAT molecule (see Extended Data Fig.6l), *in vitro* studies also show that N-terminal truncations of AAT (1-290aa and 1-323aa) are capable of forming misfolded, oligomer-like conformations^14^ which can presumably form higher-order species (Also see later, Fig.4).

**Figure 4:**
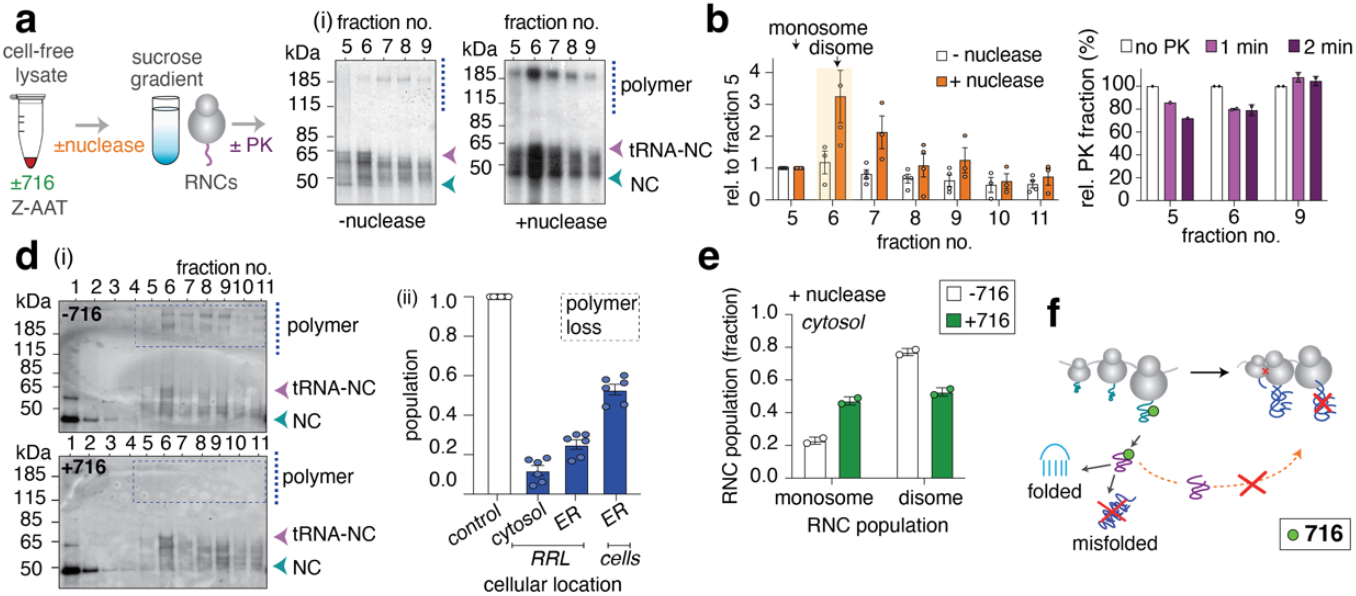
Ribosome stalling induced by co-post assembly and its modulation by a pharmacological chaperone. **(a)** Schematic image showing cell-free (RRL) expression (±716 compound) and purification of Z-AAT (after 20 min biosynthesis) via sucrose gradient. Ribosome collisions and RNC assembly formation was studied using nuclease and/or proteinase K (PK) treatment. (i) ^35^S methionine detection of selected fractions (5 to 9) before and after nuclease treatment. **(b)** Densitometry analysis quantifying the relative populations of monosomes, disomes and polysomes with and without nuclease treatment (n=4, with SEM). **(c)** Proteinase-K sensitivity of the ribosome-bound polymers in selected fractions (5, 6, 9) from the sucrose gradient (n=3, with SEM). **(d)** (i) Fractions from a sucrose gradient (^35^S methionine detection) of Z-AAT expressed in RRL (± 716). The polymer population is highlighted. (ii) Population of polymers in the presence of 716 in RRL with and without ER microsomes, and also in HEK293 cells (n=6, with SEM). **(e)** Quantification of the population of monosomes and disomes in the presence and absence of 716 in RRL (no ER microsomes), following nuclease treatment. **(f)** A schematic model for Z-AAT’s cotranslational misfolding and its modulation by the 716 compound. All gels are representative of biological repeats.

### A ribosome-bound co-translational folding intermediate drives assembly formation

Next, to provide a molecular-level understanding for how AAT RNCs participate in co-translational (mis)assembly, we used cryoEM to study purified, full-length (non-stop) wild-type AAT RNCs (no microsomes, Fig.3c & Extended Data Fig.7a). The resulting reconstruction had an average resolution of 3.7 Å (EMD-53451), while the local resolution of observable AAT NC density within the exit tunnel varied from 7 to 14 Å (peptidyl transferase centre (PTC) to the vestibule, respectively, Extended Data Fig.7a,b). A *de novo* built poly-alanine model of AAT revealed a helical turn just above the constriction site, the narrowest part of the ribosome exit tunnel. The model also indicated AAT formed a set of ribosome contacts centred around uL4, uL22 and eL39, similar to those observed in the Xbp1u stalling motif^10^, and thus likely involved in translational arrest (Fig.3c(i) & Extended Data Fig.7c). The AAT NC could be traced throughout the tunnel to the ribosomal exit. However, no NC density was observed beyond the exit tunnel, even after *in silico* 3D classification around the exit tunnel of the ribosome, likely due to the flexibility of the NC.

To overcome this challenge, we evaluated the conformational landscape accessible to the ribosome-bound, (but emerged) AAT via C-alpha, structure-based, experimentally-restrained MD simulations. Here we used the wild-type AAT-RNC cryoEM structure as a starting model (Fig.3c) and protection factors derived from a prior PEGylation analysis of RNC structure^8^ to reweight the conformational ensemble of the AAT RNC (Extended Data Figs. 8-11 and Extended Methods). We observed that ribosome-bound AAT is highly dynamic starting from the PTC and throughout the tunnel (Fig.3c(ii)), and where the ribosome-emerged NC segment is observed as an elongated ensemble of conformations (97% R_g_ ∼5.5nm vs 2.9nm for the native protein), including a population (3% total) that are more structured with native-like contacts (R_g_ ∼2.5nm Fig. 3c(iii)). The emerged segment appears largely disordered (Fig3c(iii,iv)) but has a propensity to sample native-like contacts (local Q ∼0.5) in each of the helices and some β-sheet contacts (residues T203-P255); these regions, corresponding to the B and C sheets (Fig.3c(v,vi)) & Extended Data Fig.11), are known to initiate folding in AAT^15^. Additionally, two regions (residues T180-F190 and I360-K370) showed a significant reduction in the fraction of native contacts (Q values < 0.15) and coincided with two distinct boundaries identified previously by limited proteolysis^8^ (Fig.3c(v,vi)), corresponding to N (D1-K191) and C terminal domains (G192-K365) of persistent structure. These results are also supported, in part, by biophysical and NMR studies of the N-terminal domain and show a non-native molten globule-like structure^8^.

Overall, this three-dimensional structural model suggests that AAT forms a heterogeneous ensemble of co-translational folding intermediates, including extended conformations with a propensity for secondary structure formation. This model is consistent with a molten-globule folding intermediate observed in AAT^16^ and other proteins^17^. The dynamic structural characteristics of this ribosome-bound, non-native intermediate state thus make it highly compatible for initiating both co-translational folding^8^ and assembly on the ribosome.

### Ribosome-bound assemblies are formed by recruiting released nascent species in a co-post mechanism

Next, we studied co-translational (mis)assembly pathway on the ribosome in a controlled manner by refolding partially-denatured Z-AAT NCs in the presence of RNCs (to mimic an in-cell model of NC release and assembly) (Extended Data Figs.6i,j). While no polymers were observed in the protein in the absence of RNC (Extended Data Figs.6i,j), consistent with prior polymerisation studies^18^, ribosome-bound polymers formed readily under these conditions (within 2 hours at 25°C) at relatively low concentrations (from 300nM to 1μM) in aqueous buffer (Extended Data Figs.6i,j).

We complemented these experiments by using PEGylation to follow the folding of AAT NCs released from RNCs over time (Extended Data Fig.6g,h), using the single cysteine variant, A183C, in which Cys183 is solvent in-accessible in AAT’s folded form^8^. Following its formation on the ribosome, the released Z-AAT NC typically persisted in its intermediate state (which formed cotranslationally) for several hours (Extended Data Fig.6g,h). These data thus show that the long-lived released intermediate state of Z-AAT arises at the onset of recruitment during co-translational assembly formation (Extended Data Fig.6m, “co-post” assembly). This process appears to be responsible for lowering the energetic barrier necessary to promote AAT assembly on the ribosome as shown by the vast difference in the rates at which Z-AAT polymerises on versus off the ribosome.

### Persistent ribosome pausing due to protein misfolding causes ribosome collisions

As shown earlier (Figs.2,3), translational arrest and NC recruitment are key to co-translational assembly formation, and we hypothesised that this scenario might result in ribosome collisions, which typically occur when ribosomes stall due to cellular stress. We therefore exploited the ability to produce natively-folded AAT in the absence of microsomes^8^ in an uncoupled translation reaction to study whether co-translational mis-assembly in a cytosolic environment could induce ribosome collisions (Figs.4a & Extended Data Figs.12a,b). The resulting sucrose density gradient profile revealed monosomes comprised of 2 dominant RNC sizes (∼61 and 63 kDa) and a set of decreasing sizes (Figs.4a,b & Extended Data Fig.12d). Fractions 6-11 reflected polysome formation with queued ribosomes at different stages of translation (i.e., progressively shorter lengths of the NC, Fig.4a & Extended Data Figs.12d). Ribosome-bound polymers of varied lengths were observed in the monosomes, as expected, however the polysome fractions (6-11) showed one discrete band (∼185 kDa), consistent with a ribosome-bound trimer. Next, nuclease treatment was used to differentiate between polysomes and ribosome collisions (Figs.4a,b); this resulted in a substantial collapse of the polysomes into monosomes (fractions 5) (Fig.4b), the concomitant appearance of a nuclease-resistant disome peak (fractions 6-7), and a shift in the population of the trimer band, a hallmark of pairs of collided ribosomes (Fig.4b). The polysome (fraction 9) peak (following nuclease treatment) also showed resistance to proteinase K relative to the monosome peak (Fig.4c & Extended Data Fig.12e), which suggests an alternative arrangement of ribosomes *e*.*g*., co-co assemblies in *cis* or *trans*^19^ which sequester the NC (Fig.4c). Taken together, these results show that cytosolic ribosomes translating misfolding-prone proteins can stall during co-post assembly, causing upstream collisions and high-order assemblies to form.

### A pharmacological chaperone can reverse the effects of co-translational misfolding on the ribosome

To validate the co-post assembly model, we explored whether this mechanism could be modulated using a small molecule. For this purpose, we leveraged a pharmacological chaperone, 716, which stabilises Z-AAT and prevents polymer formation in the ER of hepatocytes^20^. When cell-free reactions were pre-incubated with a fixed concentration (20μM) of 716, there was a 45% reduction in ER-bound polymers formed in microsomes (Figs.4d & Extended Data Fig.12f) and an 80% reduction in those formed in the cytosol (fractions 6-11 (Fig.4d & Extended Data Fig.12f). When tested in cells, there was a 40% reduction of ribosome-bound polymers found in the ER (Fig 4d and Extended Data Fig.12f(iii)). Additionally, as observed in RRL, there was also an enrichment of the monosome population and decrease in disome population (Fig.4e). This result shows that ribosome collisions that occur following translational arrest (due to misfolding) is reduced when co-post assembly is modulated using the drug. Combined, these results show that the co-post polymer formation on the ribosome can be rescued by modulating released Z-AAT’s kinetic folding defect. The observation of polymers in the ER even after drug treatment (albeit at a fixed concentration) reflects the resolution at which early biosynthetic events are captured: i.e., shorter NCs may be resistant to the action of this compound. The reduction in polymer formation is most consistent with 716 sequestering released Z-AAT into a conformation that prevents ribosome-associated recruitment (and with possible binding to RNCs (Extended Data Fig.12g)), which interferes with assembly with an incoming donor (Fig.4f).

### A role for the ribosome In promoting persistent pausing

Having established a link between ribosome stalling and co-translational assembly, we pursued a molecular basis for these observations by evaluating persistent pausing in a series of unrelated proteins (FLN5, firefly luciferase, TDP-43, Ras, p53, HTTex1, synuclein, transthyretin, AAT, Figs. 5b & Extended Data Figs.13a-c). Using ACTA-synchronised, coupled transcription/translation cell-free (RRL) reactions we ensured that the ribosomes go through only 1 round of translation on monosomes, and therefore express all proteins at the same level (Fig.5a & Extended Data Fig.13c). We also restricted the expression time to a short window (< 60 minutes) to account for differences in translation rates, to ensure that engagement of release factors was not rate-limiting, and to not exhaust the reaction’s energy supplies. These parameters, along with the absence of sufficient concentrations of ribosome quality control proteins and chaperones enabled us to test for persistent stalling in a protein-dependent manner. We found that several nascent polypeptides (HTTex1, AAT, TDP-43, α-synuclein, transthyretin), were retained as full-length RNCs with stalling efficiencies ranging from 40% (HTTex1 Q46, AAT) to 80% (p53 and TDP-43) while some proteins did not show stalling (FLN5, HRas, luciferase) (Extended Data Fig.13c). In AAT at least, stalling was observed on both eukaryotic (80S) as well as *E*.*coli* (70S) ribosomes (Extended Data Fig 13c).

**Figure 5:**
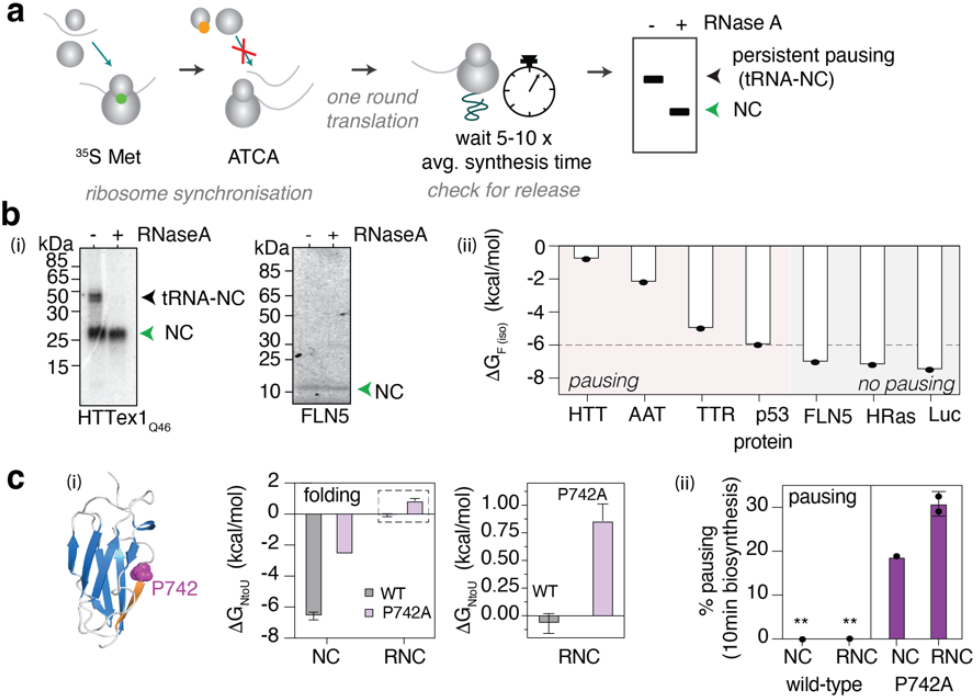
An energetic basis for persistent translational pausing for co-translational (mis)folding. **(a)** An assay for studying persistent translational pausing in cell-free (RRL) using synchronised ribosomes which undergo one round of translation. **(b)** (i) ^35^S methionine detection of persistent pausing (HTTex1) and no pausing (FLN5) (n=3, representative gel). (ii) Observed translational pausing across a series of proteins relative to their known Gibbs free energies of folding^25,36-41^ **(c)** (i) Gibbs free energy of folding of FLN5 and its destabilised variant, P742A^24^. (ii) Persistent translational pausing observed in wild-type FLN5 and the P742A variant on and off the ribosome (n=2, with SEM). All gels are representative of biological repeats.

These results suggest that a short time window exists for translational pausing at or near stop codons prior to translational termination (*ca*. 0.25s^-1^ in eukaryotes^21^). The proteins tested showed no similarity in topology, hydrophobic stretches or extent of disorder or mRNA structure around the pause site. Aside from known amino acid motifs^22^ that cause pausing (e.g., polyproline stretch in HTTex1 Fig.13b), no obvious consensus stalling sequence was observed in these polypeptides (Extended Data Fig.13b); the latter finding is consistent with comprehensive organism-wide ribosome profiling studies^23^.

One apparent striking difference between the proteins is their intrinsic thermodynamic stabilities (Gibbs free energy of folding, ΔG_N_→_U_, off the ribosome (ΔG_F_), Fig.5b(i); the non-pausing candidates typically possess larger ΔG_F_ values (> 6 kcal/mol) relative to the pausing candidates (Fig.5b(i)). We thus hypothesised that the thermodynamic stability of NCs may be a contributing factor. We tested this idea by decreasing the stability of the FLN5, an Ig-like protein which folds under thermodynamic control, and has also been used extensively to examine basic co-translational folding phenomena^9,24,25^. We introduced the P742A mutation, which destabilises the native state of FLN5 from -6.6 kcal/mol to -2.6 kcal/mol^24^. When separated by a 31 amino acid linker to the ribosome, the native state of ribosome-bound P724A is destabilised further to +0.9 kcal/mol, while the wild-type it is -0.07 kcal/mol^24^. Thus, relative to the wild-type, the variant is less able to adopt its native state at this RNC length (Fig.5c(i)). We found persistent stalling both in FLN5 P742A (∼20% of species), and when it possessed a 31aa linker (FLN5^P742A^+31, ∼30%), the latter of which ensures that the FLN5 domain has fully exited the tunnel^9^ (Fig.5c(iii)). These results revealed that destabilising point mutations can promote (persistent) translational pausing in folding-competent proteins, likely by destabilising ribosome-bound co-translational NC folding intermediates and the native state^8,24^; in turn this likely delays intrinsic NC folding rates during biosynthesis, including contributions from pulling forces generated by structure acquisition^26^. Proteins whose folding is under kinetic control or those that complete folding off the ribosome are, therefore, more likely to be affected by perturbations from point mutations and thus enhance pausing of fully-synthesised (but ribosome-bound) proteins, as modelled here by a simple NC system. These results show that the intrinsic thermodynamic (and kinetic) stability of the emerging NC also contributes to persistent ribosomal pausing.

## Discussion

Polypeptide chains are subject to competing events: protein folding and misfolding. While numerous reports have detailed co-translational folding phenomena, few studies have explored co-translational misfolding. We therefore set out to elucidate the physical basis for co-translational misfolding by creating cellular and *in vitro* snapshots of active protein biosynthesis occurring on ribosomes. This new approach enabled us to detect ribosome stalling and early newly-released protein species including kinetically-trapped intermediates and oligomers, which are hallmarks of protein misfolding. Using a pathological variant of AAT we captured direct structural and biochemical evidence for a new mode of protein misfolding and mis-assembly occurring on translating ribosomes. The mechanism underlying this process involves assembly of released polypeptides in a non-native conformation, onto ribosome-bound nascent chains also in a non-native conformation to form a structured assembly (‘co-post’ assembly), which we have shown occurs at two major cellular sites of protein folding: the ER and the cytosol (Fig.6). This assembly process is driven by the availability of kinetically trapped, partially folded protein monomers. These species can be recruited by the ribosome via interactions that they make with the molten globule characteristics of the tethered NC (Fig.6), as illustrated by the structure of a ribosome-bound Z-AAT co-translational folding intermediate. Our data thus show that protein misfolding and assembly is not exclusively limited to the post-translational self-association of mature proteins.

**Figure 6:**
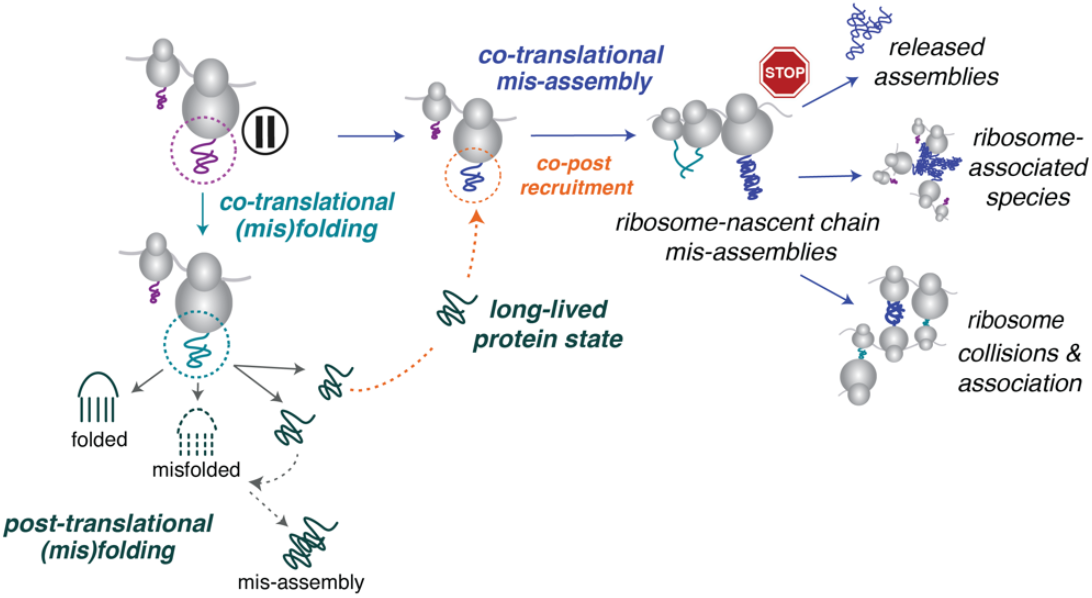
Parallel co- and post-translational protein misfolding pathways. A mechanistic model illustrating the co-existence of co- and post-translational folding and misfolding pathways for proteins.

Protein biosynthesis and the competing processes of co-translational protein folding and misfolding (Fig.6) are thus entirely coupled processes. These results highlight how any modulations in translational efficiency, mRNA defects, ribosome pausing, stalling, and cotranslational misfolding present issues that are constantly managed by the cell. Indeed, the mere existence of a co-post mis-assembly process reveals that achieving protein folding is even more perilous than first thought, particularly in the context of multi-domain and proteins acquiring quaternary structure: indeed, achieving successful folding constantly relies on an intertwined set of protein and ribosome quality control mechanisms. By extension, co-translational assembly formation, which underpins how some protein complexes may form intracellularly^27^, must similarly involve an extended ribosome-bound nascent chain capable of adopting persistent tertiary structure upon binding.

Our study also demonstrates for the first time that folding-competent, but destabilised NCs bound to the ribosome also promote stalling. This insight uncovers a new mechanism involving a polypeptide chain’s capacity to fold on the ribosome, that sits alongside known features that lead to ribosome stalling^28^, including defective mRNA^29^, amino acid motifs (e.g. proline^22^ or arginine^30^), and codon optimality at the PTC^31^. The consequences of stalled ribosomes include ribosome-associated aggregation and the release of truncated NCs; the release of such NCs may be detrimental, for example, in neurodegenerative disorders including Parkinson’s^32^ and Huntington’s^33^ diseases. A molecular basis for ribosome stalling, which is central to co-post assembly and collisions, lies in the thermodynamic (and kinetic) stability of a growing NC (i.e., the folding status of the NC). As shown with a small immunoglobulin protein (FLN5), a NC destabilised by mutation is more likely to pause on the ribosome and adversely affect the overall entropic freedom for folding^24^ than the wild-type polypeptide; point mutations thus impact on the earliest stages of folding despite tempering/buffer effects of the ribosome^8,24^. As a result, the tethered NC may be vulnerable to recruitment, should a suitable structurally compatible species also be available (Fig.6). This scenario is thus likely to extend to other misfolding-prone systems that also show persistent pausing on the ribosome, such as huntingtin (HTT) (Extended Data Fig.13 and Ref^34^) which readily oligomerises and aggregates within cells^35^. Additionally, this is a mechanism that likely underpins (functional) co-translational assembly to form homo- and heteromeric complexes (e.g. p53, Extended Data Fig.13).

As shown in detail for AAT, co-post (mis)assembly involves structural transitions and sets of interactions between a “donor” and “acceptor” that are unique relative to the released protein, since there are physical constraints imposed by the presence of the ribosome and the extent and availability of NC sequences to engage (e.g., NC lengths > 200 amino acids which expose the partially-folded N-terminal domain^8^ (Extended Data Fig.6k)). The physical constraints of the cell environment are likely to play a role: by considering the expected steric environment of the ER, we hypothesise that a ribosome-bound polymerisation mechanism could allow a more substantial C-terminal swap (*∼* 65aa) with a newly-released Z-AAT, relative to that observed for the mature protein^5^ (Extended Data Fig.6l). Critically, we found that co-translational AAT misfolding and polymerisation can also be rescued with the pharmacological chaperone, 716, illustrating that AAT polymerisation occurs co-translationally and post-translationally. The drug also revealed that by reducing polymerisation, the extent of stalled ribosomes and collisions can also be reduced. These results have important implications for protein misfolding and aggregation in other proteins, and how therapeutics exploiting early biosynthesis may serve as a potential mode of early intervention.

The approach we have developed will allow future analyses of the relative impact of co-versus post-translational misfolding in creating imbalances in proteostasis. As illustrated in the differences between the on and off ribosome rates of AAT polymerisation, co-translational misfolding and mis-assembly pathways present ‘shortcuts’ that initiate protein misfolding events: rapid transitions occurring between partially-folded RNCs and a released donor species reduce the (typically) hours timescale needed for proteins to misfold and aggregate to the timescale of biosynthesis (minutes) observed co-translationally on ribosomes. In stressed cells which fail to eliminate extremely early pathological species, it is possible that co-translational misfolding may actually precede post-translational misfolding.

In conclusion, co- and post-translational misfolding and thus aggregation, co-exist as parallel pathways as shown for two representative disease-causing proteins. Co-translational misfolding and mis-assembly on the ribosome also represent the earliest forms of NC-mediated translational dysregulation, though these species have not previously been detected in cells. This early aberrant folding and mis-assembly likely initiates a cascade of misfolding-related phenomena and interacts with the integrated stress response. Overall, our data suggest a direct relationship exists between co-translational misfolding on the ribosome and conformational disease, which has implications for describing disease pathogenesis and points to new therapeutic directions.

## Acknowledgements

We thank current and past members of the Christodoulou and Cabrita groups for useful discussions. We thank Drs. Janet Kumita and Luisa Capabello (Department of Pharmacology, University of Cambridge) for the HeLa-Trex cell line. We acknowledge the assistance of Reuben Asher for pilot studies of α-synuclein (UCL Alumni Undergraduate Summer Studentship 2023), the Erasmus+ Programme initiative (Nicoletta Carfi, Francesco Minieri), and Lincoln College, Oxford College EPA Summer Research Placement grant (Thaleesha Iynkaran). We thank Drs. Andrew Osborne and Mina Edwards (ISMB, UCL) for advice and training in cell culture, Drs. Virginia Silio and Mike Redd at the UCL-Zeiss Multiscale Imaging Centre for training in confocal microscopy, Dr. Thomas Blacker (ISMB, UCL) for guidance with fluorescence image analysis, and Dr. Adolfo Quinones Lombrana in the Darwin Research Facility for his technical advice. We acknowledge the use of UCL/Birkbeck’s EM facility (202679/Z/16/Z and 206166/Z/17/Z) and thank Drs. Shu Chen and Natasha Lukaynova for their help with electron microscopy and Dr. David Houldershaw for his help with computing and data processing. We acknowledge Diamond for access and support of the cryo-EM facilities at the UK National Electron Bio-Imaging Centre (eBIC), proposal bi20287, funded by the Wellcome Trust, the Medical Research Council and the BBSRC. We acknowledge the use of high-performance computing (HPC) at UCL (Kathleen, Young and Myriad), the UK Materials and Molecular Modelling Hub which is part-funded by the EPSRC (EP/T022213/1, EP/W032260/1 and EP/P020194/1), and the Baskerville Tier 2 HPC service (Advanced Research Computing, University of Birmingham) which is funded by the EPSRC and UKRI through the World Class Labs scheme (EP/T022221/1) and the Digital Research Infrastructure programme (EP/W032244/1). J.C. is supported by Wellcome Trust grant No. 206166/Z/17/Z. L.D.C is supported by an AlphaOne Foundation grant.

## Author Contributions

The original concept was conceived by L.D.C. and the study’s design was developed by E.P., L.P.C., J.C., and L.D.C. Data acquisition and analysis was undertaken by E.P., L.P.C., C.R.H., and L.D.C. Data interpretation was undertaken by E.P., L.P.C., C.R.H., J.O.S., A.N.M., S.H.S.C., J.C., and L.D.C. J.A.I. and D.A.L. provided the 716 compound and purified AAT polymers. Funding for the project was acquired by J.C. and L.D.C. Project administration and supervision was undertaken by J.C. and L.D.C. The manuscript was written by E.P. and L.D.C. with editing undertaken by C.R.H., J.O.S., A.N.M., S.H.S.C., J.A.I., D.A.L., and J.C.

## Experimental Procedures

### See Extended Methods for more details DNA construct design

For mammalian cell expression of RNCs the CMV-driven vector, pTREX (Invitrogen) was used. All DNA constructs used for this study were synthesised by GeneWiz. To isolate ribosome-nascent chain complexes from HEK293-Trex cells, translational arrest was induced with a modified form of the stalling motif XBp1u^10^. To produce AAT RNCs in *E*.*coli*^42^ the arrest-enhanced form of SecM (SecM-AE1)^25^ was used.

### Mammalian cell culture

T-Rex HEK293 cells grown in either 12-well, 6-well or 15cm^2^ dishes were cultured in DMEM containing blasticidin (5μg/ml) and 10 % (v/v) Fetal Bovine Serum (Tet-approved) at 5% CO^2^ in a humidified incubator. Following passaging, cells were grown to 70-90% confluency, and DNA was transfected with PEI. The cells were serum-starved for 16-18 hours (to minimise DNA fragmentation^43^ and to synchronise cell growth at G1 arrest^44^). Protein expression was induced with 1μg/ml doxycycline. For ribosome synchronisation experiments, following serum-starvation, the cells were incubated in met-free RPMI 1640 media with 10% dialysed FBS for 45 min to 1 hour followed by induction of transcription with doxycycline for 2 hours^45^. Translation was induced with 2mM L-methionine. To generate one-round of protein synthesis the cells and ribosomes were synchronised and protein expression was induced: Harringtonine was added after 3 minutes of doxycycline-induced expression^46^, for 2-5 minutes (following an approximate elongate rate of 3-5 amino acids per second^47^). For generating cell stress, 0.05μg/ml tunicamycin and 10μM of MG-132 was used. In all cases, protein translation was quenched with 100μg/ml cycloheximide for 5 minutes prior to cell harvesting. Cells were immediately flash frozen in liquid nitrogen prior to analysis.

### Immunocytochemistry and confocal scanning microscopy

Cells (70-80% confluency) were seeded on No.1.5 cover slips and RNCs were expressed for 5 and 48 hours. Cells were fixed with 4% w/v paraformaldehyde for 30mins, permeabilised with Triton X-100, and probed with 0.5μg/ml of AAT-specific antibodies, 1C2 (total AAT) and 2C1 (AAT polymers), and 5μg/ml of the following antibodies: rabbit anti-RPS10 Alexa 697-conjugated, and goat anti-Sec61α. Secondary antibodies (Alexa 488 and Alexa 555) were used at a concentration 1 μg/ml. Nuclei were stained with 1μg/ml DAPI (Cambridge Bioscience). Cells were mounted using VectaShield Anti-Fade on #1.5 coverslips and sealed with clear varnish. The slides were visualised using a Zeiss AiryScan 980Vis microscope at 63x with Immersol immersion oil. Excitation and emission windows were set to minimise spectral overlap observed between the closely-related fluorophores.

### Manders correlational coefficient analysis

Confocal and super-resolution data (Airyscan) were collected and analysed using Fiji software. Images were filtered and a threshold was applied to reduce high frequency noise. Mander correlation coefficients, M1 and M2, were calculated using Coloc2 plugin. A standard error of the mean was taken from 4 measurements, and where necessary, a one-tailed T test was applied to calculate differences in fluorescence intensities observed over time.

### Isolation of cytosolic and ER-bound RNCs from mammalian cells

Cell pellets were resuspended in lysis buffer (20mM HEPES pH 7.8, 6mM MgCl_2_, and 100mM KCl, 1mM DTT, 10% protease inhibitors, 0.35 mg/mL cycloheximide) and lysed using homogenizer (Isobiotec) with 8mm diameter ball. The lysates were spun down at 600xg for 5min at 4°C twice to remove cellular debris and 1,200xg for 5 min to remove the plasma membrane. The supernatant was then spun down at 20,000xg for 35 min to remove any plasma membrane contamination. The resulting supernatant was then either spun at 100,000xg for 60 min to isolate the crude ER fraction (pellet) from the cytosol (supernatant), or alternatively, loaded onto a 20-60 % (w/v) sucrose gradient in lysis buffer spun at 55,000 rpm for 60 min (TLS-55 rotor, Optima Max centrifuge Beckman Coulter) to separate the ER-bound RNCs from any cytosolic RNCs and other proteins. 1.2 mL sucrose gradients were hand-poured by layering increasing concentrations of sucrose into centrifuge tubes. The gradients were incubated on ice for 1-2 hours on ice prior to the application of sample. Fractions were collected manually from the top of the tube and subjected to PAGE followed by immunoblotting.

### Cell-free protein synthesis and PAGE analysis of RNCs and NCs

Cell-free, coupled transcription-translation, translation-only, and aurintricarboxylic acid (ATCA) synchronised reactions were carried out in rabbit reticulocyte lysate (RRL) (with and without canine pancreatic microsomes) (Promega) as described previously^8. 35^S methionine and fluorographic-enhanced autoradioagraphy (Enlightening, Revvity) was used for detection^8^. RNC samples were separated on 8% w/v Bis-Tris pH 7.0 gels, using a pH 5.7 loading dye to preserve the tRNA-bound nascent chain. NCs were released using 1mg/ml RNase A. Samples were deglycosylated with PGNaseF using the manufacturer’s instructions. For the detection of polymers by 2C1, a non-denaturing loading dye was used to preserve the integrity of the polymer species. Densitometry of gels was undertaken using Fiji and Fityx as previously described^8^ .

### Immunoblotting (Western blot)

Following PAGE, samples were transferred at 100V for 1 h at room temperature onto nitrocellulose membrane. Membranes were blocked for 1 hour in 5% skim milk powder in TBS with 0.1% w/v Tween-20. *See Supplementary Table 1* for all antibody concentrations. Primary antibodies were incubated for a minimum of 1 hour at room temperature or up to 18 hours at 4°C, followed by incubation with secondary antibodies in in TBS with 0.1% Tween-20 (1 hour, room temperature) and detection was with SuperSignal West Pico or Femto chemiluminescent substrate (ThermoFisher) and imaged using Odyssey Fc Imager (LICOR).

### Negative-stain electron microscopy of RNC assemblies

The samples were diluted with EM buffer (50 mM HEPES, 100 mM KOAc, 5 mM Mg(OAc)_2_, 0.5 mM TCEP, pH 7.4) to a final ribosome concentration of 30 nM. The diluted samples were adsorbed onto carbon coated grids (CF-300 Cu, EMS), freshly glow-discharged at 5x10-5 Torr, 35 mA, for 90 sec. After 90 sec of incubation, the excess solution was blotted and the grid was washed twice by floatation on top of 80μL droplets of EM buffer. The sample was then stained with 2% w/v uranyl acetate, by floating the grid immediately on the surface of the staining solution for 50 sec. After removing the excess solution with blotting paper, the grids were examined onto a Technai T12 at 67,000x magnification. AAT polymer lengths were calculated in Image J using a method described previously^48^.

## Notes

### Competing Interest Statement

The authors have declared no competing interest.

