## Extended Data for "Visualisation of translating ribosomes reveals the earliest steps of protein misfolding in human disease"

**Extended Methods**

*Expression and purification of AAT 70S RNCs and isolated AAT from E.coli*
AAT RNCs were expressed and purified from *E.coli* as described previously[42,49]. AAT was purified on AAT-Select resin as described previously[8]

*Seeded aggregation studies*
For the investigation of co-post assemblies in buffer, Z-AAT DNA constructs containing a stop-codon were expressed in RRL for 90 min. The reaction was purified via Ni-IDA affinity chromatography under completely denaturing conditions (8M urea) to purify Z-AAT in its unfolded state (reaction 1). Z-AAT RNCs were produced using the non-stop codon linear DNA. The reaction was quenched with 1mg/ml emetine, the RNCs were purified via a 30% (w/v) sucrose cushion and the pellet was resuspended in RNC buffer (reaction 2). Reaction 1 was then mixed with reaction 2, resulting in a 0M effective urea concentration in the final mix. The final reaction mix was then incubated at 25$^{o}$C for 120 min and aliquots were withdrawn and analysed by semi-denaturing PAGE.

*PEGylation of released AAT over time*
PEGylation reactions were undertaken as described previously[8]. Briefly, A183C/C232S AAT RNCs were generated in RRL and purified by a sucrose cushion. The release of the NCs was induced with 1mg/ml RNase A and the reactions quenched with 1mM PEG-5K maleimide. The samples were run on a Laemmli gel.

*mRNA cell-free expression & analysis of ribosome collisions via micrococcal nuclease and proteinase K*
Z-AAT mRNA with stop codon was produced from linear DNA using HiScribe T7 ARCA mRNA kit (with tailing) (NEB). Z-AAT (using 100ng/µL mRNA as a template) was generated by *in vitro* translation using nuclease-treated RRL (Promega), at 30°C for 30 min. The reaction was quenched with 1mg/ml emetine and applied to a 1.2mL, 20-60% (w/v) sucrose gradient. The samples were spun in an Optima MAX centrifuge, using a TLS-55 rotor at 55,000 rpm, for 1h at 4°C, with the slowest acceleration and deceleration settings. Fractions were collected manually and polysome profiles were generated to assess the nature of the ribosomes (monosome vs polysome). For polysome dissociation each 40 µL reaction was treated with 0.2 units of micrococcal nuclease and supplemented with 3mM $CaCl_2$. The reactions were incubated at 28$^{o}$C for 2 min and nuclease activity was then quenched with 6mM EGTA. The resulting samples were then applied to a 20-60% (w/v) sucrose gradient as described above.

*Assessing persistent translational pausing*
DNA constructs (with a stop codon) corresponding to each protein of interest was expressed in a coupled transcription-translation in RRL. RRL is typically limited in the concentrations of molecular chaperones and ribosome quality control (RQC). The coupled reactions were used to exclusively generate monosomes and avoid pausing due to ribosomes collisions formed within polysomes. For ribosome synchronisation and one round of translation, protein biosynthesis was initiated with $^{35}$S methionine, followed by the addition of 75µM ATCA after 45s. Note for Z-AAT expressed on 70S ribosomes (Extended Data Fig.13c), the PURexpress system (NEB) was used. The expression time window for each construct was set for 5 to 10 times the average time needed to generate the protein (assuming a translation rate of 1 amino acid/second). RNCs observed following this time window were deemed to be the result of persistent pausing. The choice of the short time window imposed was due to the following: (i) to ensure that the components of the RRL reactions were not exhausted; (ii) to factor in translational kinetics and enable all ribosomes to complete one round complete of synthesis; (iii) to consider any slow kinetics associated with release factors (which may be concentration-limited within RRL).

*716 binding by fluorescence polarisation*

Fluorescently-labelled (Alexa 488) 716 (20 nM) was added to duplicate samples and the reactions were incubated for 30 min at room temperature. Reactions were then split into the wells of the plate and fluorescence polarisation was measured with 5 seconds shaking before reading.

*CryoEM of RNCs*

Glow discharged grids at 5x10-5 Torr, 35 mA, for 40 sec, were first washed inside a droplet (80 µL) of 0.2 mM N-dodecyl-β-D-maltoside diluted in water. The grid was then blotted with Whatman blotting paper until dry. 3 µL graphene oxide diluted in water, to a final concentration of 0.2 mg/ml was then applied and incubated for 1 min on the grids, before the excess solution was blotted[50]. Purified RNCs were then applied on the grid to 100 nM final concentration, diluted in EM buffer. The grids were subsequently vitrified by plunge freezing in liquid ethane using Vitrobot (FEI) and a wait time of 10 sec, blot time 14.5 sec and blot force -10 pN. Vitrobot chamber conditions were 4°C and 100% humidity. Data were collected at eBIC at Diamond (em20287-34), on a Titan Krios I microscope with 300 keV acceleration voltage and a K2 detector. 5854 movies were collected. The nominal magnification was 130x and the pixel size 1.047 Å. Pre-processing (motion correction and CTF estimation) of the micrographs was performed in Relion 3.1[51]. Micrographs with a resolution lower that 5 Å were removed from the dataset (200 micrographs). The selected 5,654 micrographs were subjected to Gautomatch picking and yielded 254,976 particles. The particles, downsized by a factor of four, were subjected to a reference-free 2D classification. After visual inspection of the 2D classes, 77,946 particles were selected and 3D refined to a reconstruction using an 80S Xbp1 RNC map from rabbit reticulocyte as reference (EMDB-4729) with low-pass filtering at 50Å. Further heterogeneity of these particles was classified. Briefly, the initial reconstruction was subjected to 3D classification yielding 3 main classes; one class with preferred orientations that demonstrated steaky electron density (41.9%), one class that demonstrated a low-resolution large subunit with sparse density of the small subunit (16.4%) and finally one class with high-resolution density map (4 Å) which at higher contour level shows density at the P-site of the PTC, indicating that is occupied by a NC (32.4%). The high-resolution class with 25,621 particles, that were re-extracted and unbinned, was subjected to further 3D classification with a mask at the inter-subunit space to specifically select for particles with P-site tRNA and thus NC occupancy. The class yielding RNCs with a P-site tRNA (3,713 particles) was refined with a 50S mask producing a 7.3 Å resolution map and after CTF refinement and particle Bayesian polishing reached a 3.7 Å resolution. The Fourier shell correlation (FSC) calculation between independently refined random half subsets and the local resolution estimation were done in RELION 3.1. The cryoEM density map of the RNC is available at the Electron Microscopy Data Bank with accession code EMD-53451. Note that the validation report indicates a resolution of 6.77 Å obtained from unmasked density map. The higher resolution (3.7 Å) is achieved with a 50S mask in RELION 3.1 to overcome the flexibility of the small ribosomal subunit, which otherwise reduces the overall map resolution.

*Molecular dynamics simulations of AAT RNCs*

Coarse-grained Molecular Dynamics Simulations with the structure-based potential generated with SMOG 2.4.5[52]. For the starting model, the structure of the large 60S subunit of the ribosome and 22 AAT nascent chain residues from the PTC (K394) to the vestibule (L373) were taken from the Z-AAT RNC cryoEM structures. A further 20 residues from the vestibule were built as an extended chain in Chimera and the rest of the AAT NC taken from the crystal structure of wild-type AAT (PDB: 1QLP). During the simulations, the ribosome atoms were frozen as well as AAT's C-terminal residue, K394, to prevent NC release. The structure-based potential of the NC was defined by the wild-type AAT crystal structure (PDB: 1QLP). The C-alpha model was used for coarse graining where each amino acid residue is represented as a bead centred on the C-alpha bead and where each RNA base is represented by 3 beads that are centred on the P, C4' and N3 atoms[49]. In the structure-based, C-alpha model, the non-local contact potential interacts via a 12-10 Lennard-Jones potential and non-local contacts were defined using the Shadow Map algorithm[52]. Electrostatics interactions were introduced between the ribosome and AAT NC beads using the Debye–Hückel theory and parameters based on previous work[8].

Parallel Tempering Metadynamics (PTMetaD) were used to more efficiently sample the free energy landscape[53]. This process involved running 20 replicas across a range of temperatures where for every 5000 steps, the MD engine attempted to exchange atomic coordinates at neighbouring temperatures based on the Metropolis criterion[54]. For the structure-based simulations set up in SMOG, reduced units were applied with length, time, mass and energy scale set to 1, except for the Boltzmann constant ($k_B$) which was set to 0.00831451 ($kJmol^{-1}$ which is default in GROMACS). The use of reduced units means that there is no direct link between timescales and temperature in simulation to experiment. Therefore, simulation temperatures were chosen to span a range of unfolded and folded states and so that the mid-point temperature had approximate equal populations of folded and unfolded species. 20 replicas were subsequently set at temperatures 150-160K at 0.5K intervals. Metadynamics was applied using fraction of native contacts (Q) as a collective variable. Gaussians corresponding to the bias potential were applied every 2500 steps with a height of 0.7 kJ, width of 0.01 and the bias factor was equal to 10. Simulations were run for a total of $1x10^9$ steps for each replica at a timestep of 0.0005 ps using the stochastic dynamics integrator and time constant for temperature coupling was set to 1 ps. Frames were saved every 10 ps with the first 20% of the trajectories being discarded for equilibration. Simulations were run using GROMACS 2019.3[55] using Plumed[56] to implement metadynamics. To estimate free energy and construct the free energy landscape sampled during the simulations, reweighting of the ensembles was completed using the method described by Ref[57]. Back-mapping of coarse-grained structures for an all-atom analysis and visualisation was undertaken using cg2all software[58] with a following refinement of sidechains and clash removal using vacuum energy minimisation in the CHARMM36m force field[59].

*Ensemble reweighting using PEGylation data*

The ensemble was refined using experimental data derived from a prior study of AAT RNCs[8] which used PEGylation as measure of solvent accessibility using single cysteines as reporters. The predicted protection factors derived from the rates of PEGylation of 12 single cysteine variants in the wild-type AAT RNC were used for ensemble refinement[8]. The ensemble protection factor for each of the 12 cysteine sites were predicted using a simplification of the phenomenological approximation which is used to calculate protection factors ($P_i^{sim}$) from amide hydrogen/deuterium exchange[60] (Equation 1)

$$lnP_i^{sim} = \alpha_c N_{c,i} + \alpha_h N_{h,i} \quad \text{(Equation 1)}$$

Where $N_{c,i}$ is the number of contacts of residue *i* to other residues and $N_{h,i}$ is the number of hydrogen bonds formed by the amide hydrogen of residue *i* and $\alpha_c$ and $\alpha_h$ are the respective weights. Since protection from PEGylation is expected to be only related to burial of a residue, for comparison to protection factors derived from PEGylation rates ($P_i^{exp}$) this approximation can be simplified to (Equation 2):

$$lnP_i^{sim} = \alpha_c N_{c,i} \quad \text{(Equation 2)}$$

We use the relationship $lnP_i^{sim} = lnP_i^{exp}$ to compare the ensemble-predicted protection factors to those that had been previously derived experimentally. The number of contacts, $N_{c,i}$, was calculated for each cysteine site by calculating the number of calpha beads within a radius, $r_0$, and excluding neighbouring beads, $nb$, this was averaged across the ensemble. A C-alpha structure-based model (SBM) of wild-type AAT was used to generate a native state ensemble which we used to fit $r_0$ and $nb$ to the experimental protection factors from PEGylation rates for isolated wild-type AAT ($P_i^{exp}$). To also ensure local chain interactions were not contributing to the predicted protection factor, $r_0$ and $nb$ were fit against a random coil-like model. This was generated by removing the non-local contact interactions from the C-alpha SBM of AAT and fitting $lnP_i^{exp} \approx 0$ . The lowest RMSE to both native state and random coil was obtained with $r_0 = 1.4nm$ and a $nb \pm 8$ residues from the cysteine site bead was used (Extended Data Fig.8a,b). $\alpha_c$ was also fit to the experimental protection factors of isolated wild-type AAT and found $\alpha_c = 0.273$ (Extended Data Fig.8c,d). The simulations do not effectively sample loop dynamics and therefore the protection factor predictions of the c-terminal labelling sites (350C/355C/360C) which correspond to the reactive centre loop were not included in the fitting for the forward model.

The forward model together with the inclusion of the ribosome beads (for the calculation of $N_{c,i}$). was then used to generate the RNC ensembles. For each replica from the PTMetaD simulations, the $lnP_i^{sim}$ for the 12 cysteine sites was predicted and an RMSE to the experimental protection factors from PEGylation rates of wild-type AAT RN ($P_i^{exp}$) was used to select the ensemble which best agreed to the experimental data and this was used for ensemble for reweighting (Extended Data Fig.9).

The C-alpha SBM AAT RNC ensemble was refined by reweighting the ensemble against the protection factors derived from PEGylation rates from the wild-type AAT RNC using the Bayesian Inference of Ensembles (BioEn) software and method[61]. The L-curve analysis to select the optimal $\theta$ value by plotting the $S_{KL}$(entropy) on the x-axis and the reduced $\chi^2$ value (goodness of fit) on the y-axis for a series of $\theta$ values and selecting the elbow (θ)[61] (Extended Data Fig.10).

*Ensemble structural analysis*

Plumed[8] was used to calculate (Equation 5) a per-residue fraction of native contacts (Q) where[62]:

$$Q(X) = \frac{1}{N}\sum_{ij}\frac{1}{1+e^{\left(\beta\left(r_{i,j}-\lambda r_{i,j}^{0}\right)\right)}} \quad \text{(Equation 5)}$$

Where for $N$ pairs of contacts $(i,j)$, $r_{i,j}$ and $r_{i,j}^{0}$ are the distances between atoms in frame $X$ and the template structure (PDB: 1QLP). $\beta$ is a smoothing parameter (5Å$^{-1}$) and $\lambda$ accounts for the fluctuations of contacts and is set at 1.5. For residue Q, the calculation of Q only involved pairs of contacts for a given residue and these contacts are defined by the shadow map algorithm in SMOG[52]. Residues with only 1 contact pair predicted were not included in calculations for residue Q

Secondary structure was calculated on the back-mapped, all-atom ensemble of the ribosome-bound nascent chain using the GROMACS DSSP module, which uses the DSSP algorithm, and which detects specific hydrogen bond patterns to identify secondary structure features[63]. Visualisation and distance measurements of the top weighted structures were undertaken in ChimeraX[64].

**DNA sequence used in this study**

**Z-AAT with N-terminal signal sequence and arrest-enhanced Xbp1u**
ATGCCGTCTTCTGTCTCGTGGGGCATCCTCCTGCTGGCAGGCCTGTGCTGCCTGGTCCCTG
TCTCCCTGGCTGAGGATCCCCAGGGAGATGCTGCCCAGAAGACAGATACATCCCACCATGA
TCAGGATCACCCAACCTTCAACAAGATCACCCCCAACCTGGCTGAGTTCGCCTTCAGCCTAT
ACCGCCAGCTGGCACACCAGTCCAACAGCACCAATATCTTCTTCTCCCCAGTGAGCATCGC
TACAGCCTTTGCAATGCTCTCCCTGGGGACCAAGGCTGACACTCACGATGAAATCCTGGAG
GGCCTGAATTTCAACCTCACGGAGATTCCGGAGGCTCAGATCCATGAAGGCTTCCAGGAAC
TCCTCCGTACCCTCAACCAGCCAGACAGCCAGCTCCAGCTGACCACCGGCAATGGCCTGTT
CCTCAGCGAGGGCCTGAAGCTAGTGGATAAATTTTTGGAGGATGTTAAAAAGTTGTACCACT
CAGAAGCCTTCACTGTCAACTTCTGGGACACCGAAGAGGCCAAGAAACAGATCAACGATTA
CGTGGAGAAGGGTACTCAAGGGAAAATTGTGGATTTGGTCAAGGAGCTTGACAGAGACACA
GTTTTTGCTCTGGTGAATTACATCTTCTTTAAAGGCAAATGGGAGAGACCCTTTGAAGTCAA
GGACACCGAGGAAGAGGACTTCCACGTGGACCAGGCGACCACCGTGAAGGTGCCTATGAT
GAAGCGTTTAGGCATGTTTAACATCCAGCACTGTAAGAAGCTGTCCAGCTGGGTGCTGCTG
ATGAAATACCTGGGCAATGCCACCGCCATCTTCTTCCTGCCTGATGAGGGGAAACTACAGC
ACCTGGAAAATGAACTCACCCACGATATCATCACCAAGTTCCTGGAAAATGAAGACAGAAGG
TCTGCCAGCTTACATTTACCCAAACTGTCCATTACTGGAACCTATGATCTGAAGAGCGTCCT
GGGTCAACTGGGCATCACTAAGGTCTTCAGCAATGGGGCTGACCTCTCCGGGGTCACAGA
GGAGGCACCCCTGAAGCTCTCCAAGGCCGTGCATAAGGCTGTGCTGACCATCGACAAGAA
AGGGACTGAAGCTGCTGGGGCCATGTTTTTAGAGGCCATACCCATGTCTATCCCCCCCGAG
GTCAAGTTCAACAAACCCTTTGTCTTCTTAATGATTGAACAAAATACCAAGTCTCCCCTCTTC
ATGGGAAAAGTGGTGAATCCCACCCAAAAATATTATGATCCGCCGATTAACCGCTGGGGCC
GCCATAACTGCGCGTGGCATCCGAAAATGAACGGCTAATAA

**Table 1: Antibodies used in this study**

| Name | Catalogue Number | Type | Dilution/concentration used Western blot or (confocal) |
|---|---|---|---|
| 1C2 | SAB4200198 (Merck) | human antitrypsin, monoclonal | 1:1250 (5μg/ml) |
| 2C1 | HM2289 (Hycult) | human antitrypsin polymers, monoclonal | 1:2500 (5μg/ml) |
| A11 | AHB0052 (Merck) | rabbit antibody against oligomers | 1:1000 |
| anti-mouse IgG HRP-linked | 7076 (Cell signalling technology) | anti-mouse secondary | 1:5000 |
| anti-rabbit IgG HRP-linked | 7074 (Cell signalling technology) | anti-rabbit secondary | 1:5000 |
| Sec61α | EPR14379 (Abcam) | rabbit antibody against Sec61α | (5μg/ml) |
| rps10-Alexa 647 | EPR8545 (Abcam) | conjugated rabbit primary antibody | (5μg/ml) |
| calnexin | ab219644 (Abcam) | goat polyclonal to calnexin -C-terminal | 1:1000 |
| Rabbit anti-mouse IgG (H&L) Alexa Fluor 488 | ab150113 (Abcam) | Mouse secondary antibody | (1μg/ml) |
| Goat anti-rabbit IgG (H&L) Alexa Fluor 555 | ab150078 (Abcam) | Rabbit secondary antibody | (1μg/ml) |

**Table 2: Oligonucleotide sequences used in this study**

| Primer | DNA Sequence 5'-3' |
|---|---|
| T7 Forward | CTCGATCCCGCGAAATTAATACG |
| AAT RNC Reverse | TTTTTGGGTGGGATTCACCACTTT |
| AAT - Stop Reverse | GGCGAGCTCTTATTTTTGGGTGGGATT |
| Enhanced Xbp1u Forward | TATTATGATCCGCCGATTAACCGCTGGGGCCGCCATAACTGCGCGTGGCATCCGAAAATGAACGGC |

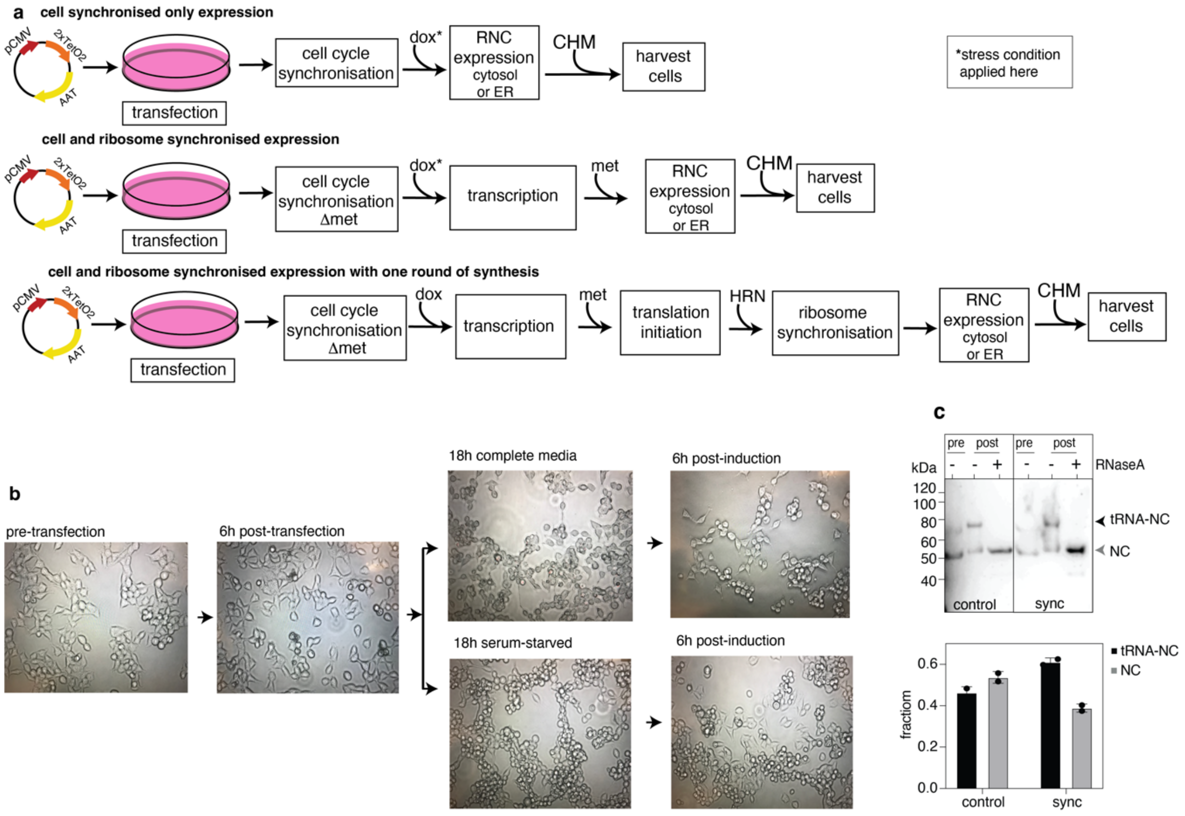


**Extended Data Figure 1: Production and characterisation of ribosome-bound polymers *in vivo* using confocal fluorescence microscopy (a)** Expression schemes used in HeLa and HEK293 cells to study AAT's early biosynthesis. **(b)** Images of HEK293 cells during different stages of transfection (Z-AAT-Xbp1u RNC) and induction with and without serum starvation to synchronise the cells. After 18 hours of serum starvation (G0/G1 arrest), typically more spherical cells are observed and used as a crude visual indicator of cell synchronisation. **(c)** *(upper)* 1C2 western blot showing expression of Z-AAT-Xbp1u with and without serum starvation to synchronise the cells. *(lower)* Quantification of RNCs and NCs from the western blot (upper). Following synchronisation, there is proportionally more RNC than NC.

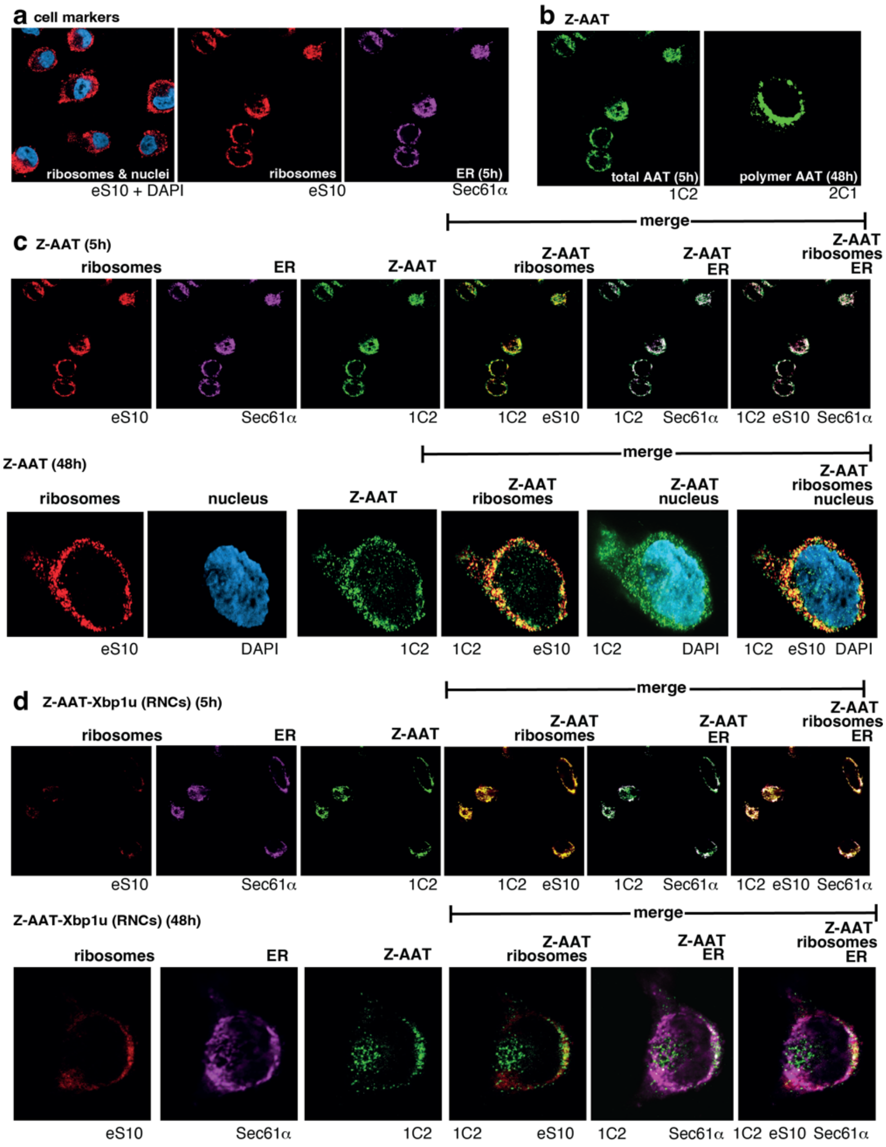


**Extended Data Figure 2: Confocal microscopy images of Z-AAT in HeLa cells (a)** Confocal images of the individual antibodies used for immunofluorescence. Also included are examples of DAPI staining of nuclei. **(b)** Detection of total Z-AAT and polymeric Z-AAT. **(c)** confocal images of isolated Z-AAT. Note that colocalisation of the ribosomes with the nucleus was used as a negative control for our image analysis; as expected, the Manders colocalisation coefficient yielded is close to zero. **(d)** Confocal fluorescence microscopy images of Xbpu1-arrested Z-AAT RNCs.

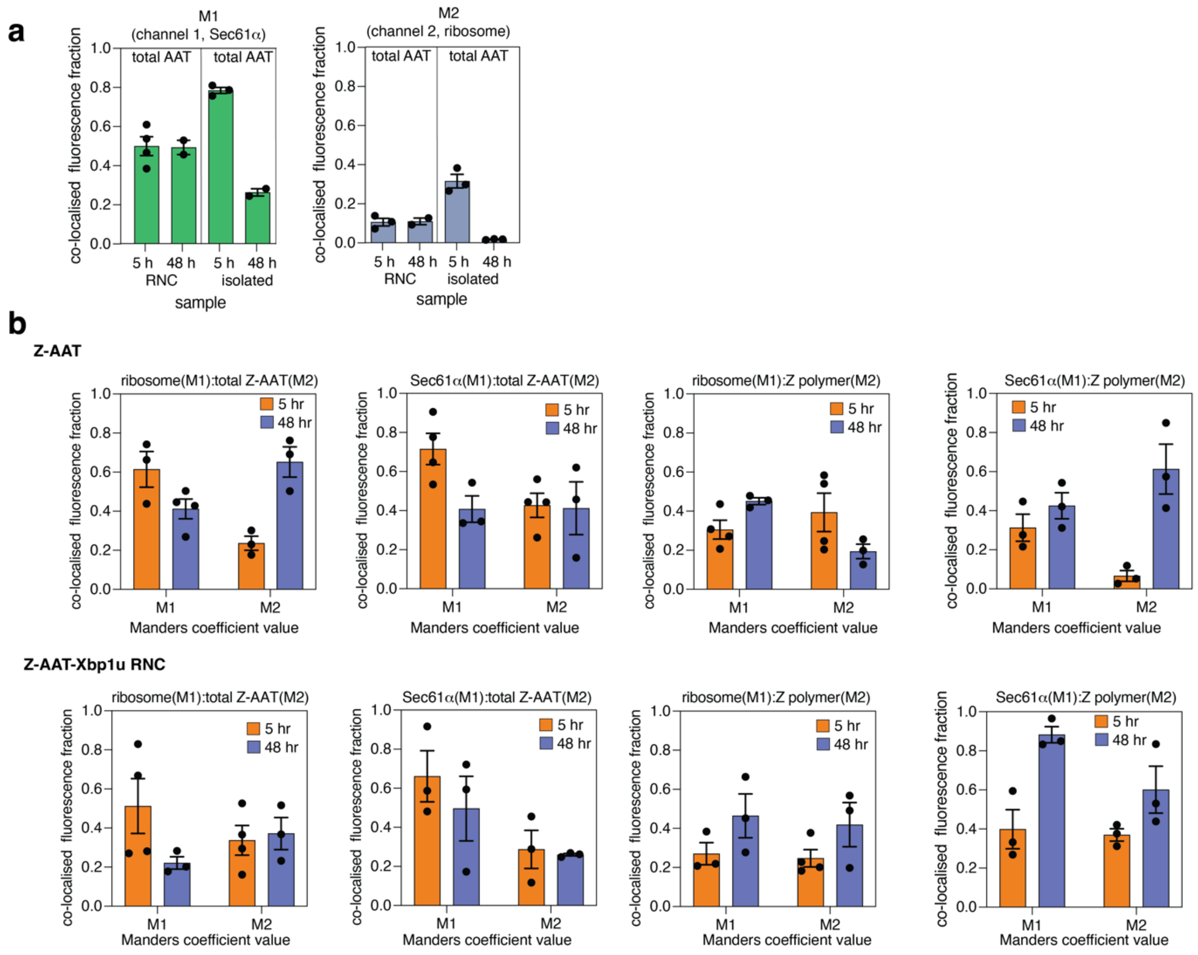


**Extended Data Figure 3: Co-localisation analysis of Z-AAT as expressed in HeLa cells (a)** Manders correlation coefficient values calculated for the fraction of colocalised fluorescence detected between the ER (channel 1) and the ribosome (channel 2) as observed across time in both isolated Z-AAT and Z-RNCs (n > 2, with SEM). **(b)** Fraction of colocalised fluorescence of Z-AAT in ribosomes and the ER as calculated using Manders correlation coefficient values. The ribosomes or ER are in channel 1 and Z-AAT is in channel 2.

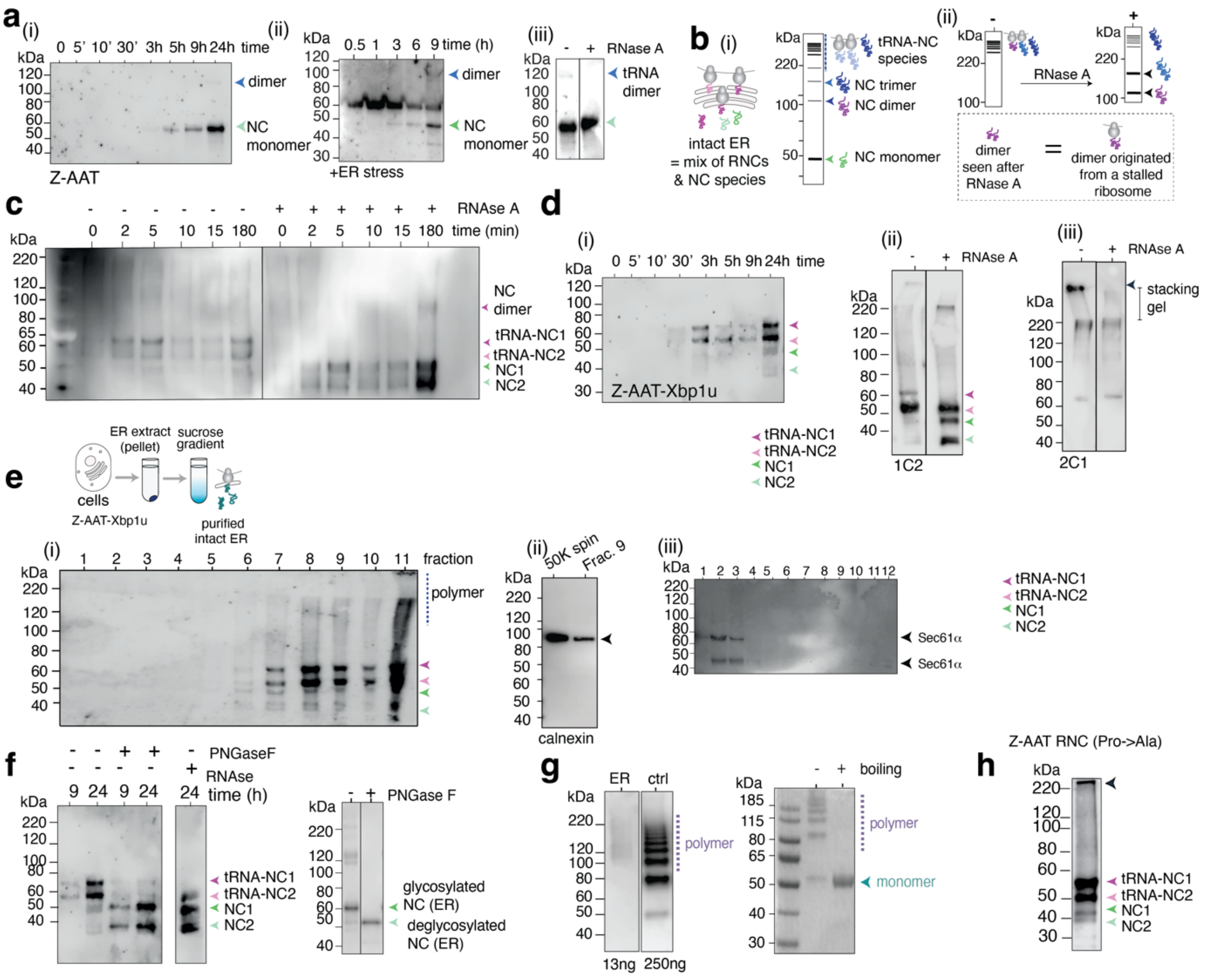


**Extended Data Figure 4: Characterisation of Z-AAT biosynthesis as produced within the ER** **(a)** (i) Time course of Z-AAT (unstressed conditions) as observed within the ER (green arrow). After 24 hours a low amount of dimer is detected (purple arrow). Note that this dimer is glycosylated and thus does not migrate at its true molecular weight on a semi-denaturing gel (compared to the stressed sample shown in (ii), which also lacks glycosylation). (ii) Time course of Z-AAT in the ER under stressed conditions in a 9-hour time period. (iii) RNAseA treatment of a sample after 24h of expression (from (i)). Note the downshift of the dimer band (blue arrow) indicating that is ribosome-bound. **(b)** (i) Schematic showing all the species observed in the gels in a(i) before and (ii) after RNAseA treatment. **(c)** Xbp1u-arrested Z-AAT RNCs (48 hours expression) with and without RNaseA treatment. **(d)** (i) Time course expression of Z-AAT-Xbp1u-RNCs in the ER (unstressed conditions). (ii) Total AAT (1C2) detection highlighting the RNC and released NC formed. (iii) Polymer (2C1) detection highlighting the presence of ribosome-bound HMW species observed in the stacking gel which are released upon RNaseA treatment. **(e)** Schematic showing the purification process of Z-AAT-Xbp1u RNCs from HEK293 cells. (i) Fractions of Z-AAT-Xbp1u RNCs taken from a 20-60% (w/v) sucrose gradient. The gradient separates the ER on the basis of its size (i.e., ER microsomes loaded with different populations of ribosomes). ER-bound AAT species were typically most abundant within heavier fractions (8-12). (ii) anti-calnexin western blot showing the presence of the ER following purification (crude ER fraction "50K spin" and in Fraction 9). (iii) anti-Sec61α western blot of fractions from a sucrose gradient of Z-AAT-Xbp1 RNCs in the ER. The detection of Sec61 was lower in heavier fractions **(f)** *(left)* PNGaseF treatment of RNCs (denaturing protocol). *(right)* PNGaseF treatment of isolated Z-AAT after 48 hours of expression. This treatment (neutral pH, with boiling) downshifts the high-molecular weight bands to monomers that are deglycosylated, suggesting that the polymers are assembled with released, glycosylated species. The RNC is also glycosylated. **(g)** *(left)* Polymers found in the crude ER fraction as detected by 2C1 under non-denaturing conditions and at neutral pH. The control (ctrl) is a sample of purified *ex vivo* polymers from Z-AAT patients' livers *(right)* Purified *ex vivo* polymers as separated on a semi-denaturing gel at pH 5.7, with and without boiling treatment. **(h)** Z-AAT-Xbp1 RNC with Pro360-Pro361 to Ala mutations. Polyprolines are a known ribosome-stalling motif[22], however two stalled species are observed suggesting that one of them is the stop codon.

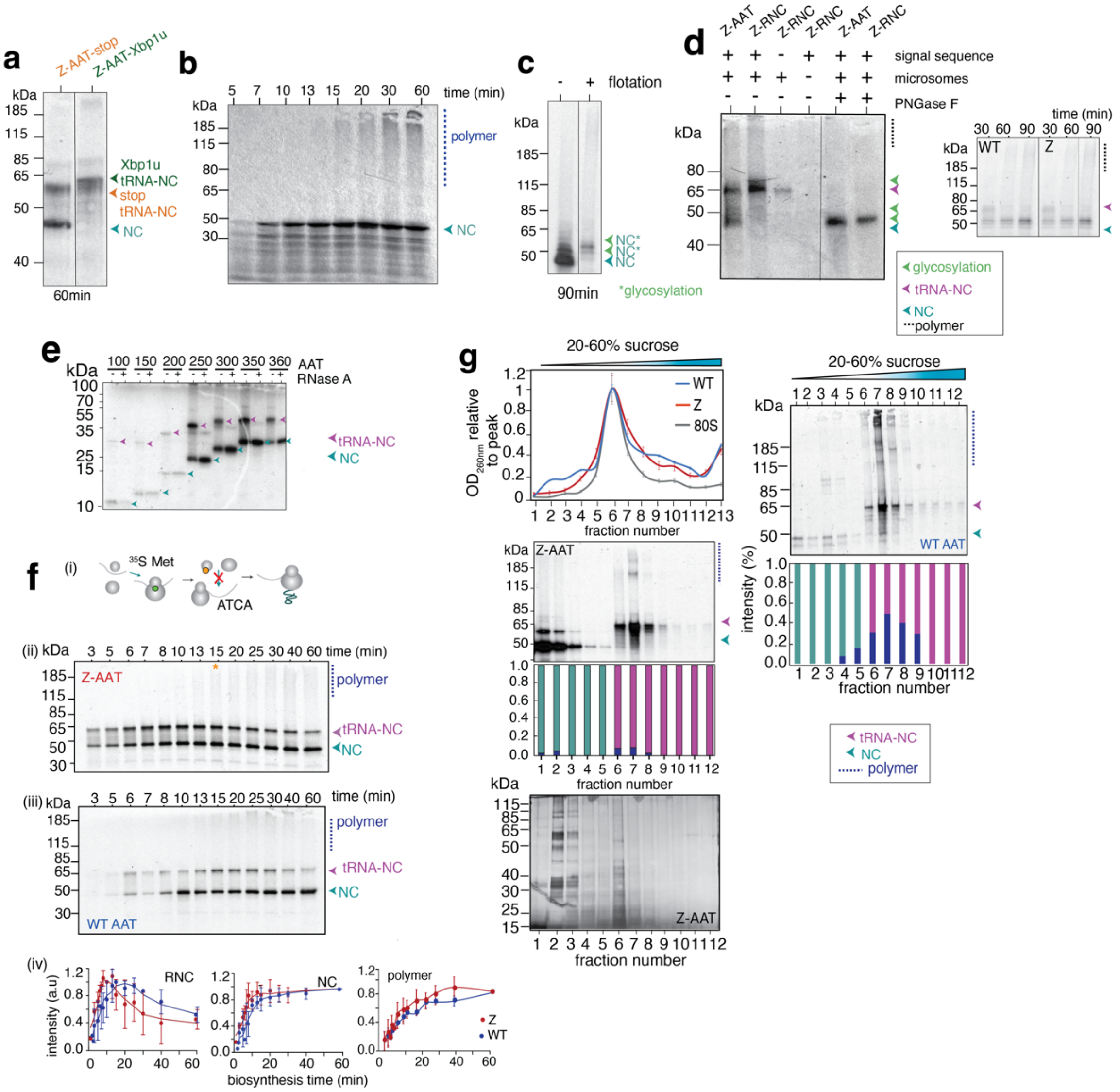


**Extended Data Figure 5: Production and characterisation of ribosome-bound polymers *in vitro*** **(a)** Expression of Xbp1u-arrested AAT in a coupled transcription-translation reaction after 60 mins. **(b)** Expression of Z-AAT in nuclease-treated RRL from mRNA template with RNaseA treatment. **(c)** Shows the enrichment of ER-bound Z-AAT following flotation **(d)** *(left)* Deglycosylation of Z-AAT RNCs with PGNaseF. *(right)* Wild-type and Z-AAT time course expression in the microsome of RRL, demonstrating natural pausing. **(e)** Synchronised reactions of C-terminal truncations of AAT with stop codon, showing spontaneous RNC formation (i.e., persistent pausing). **(f)** (i) Schematic showing the process followed to synchronise the ribosomes and express AAT in RRL. (ii) Expression profile of wild-type AAT in coupled transcription/translation RRL, separated on a semi-denaturing gel at pH 5.7. (iii) as for (ii) but for Z-AAT. (iv) Quantification of released NC, RNC and polymers formed in wild-type and Z-AAT during RRL expression. **(g)** *(left, upper)* Sucrose gradient profile (n=3, SEM), *(mid)* $^{35}$S Met detection of fractions and *(lower)* silver-stained gel of Z-AAT expressed in RRL. *(right)* Fractions recovered from a sucrose gradient from wild-type AAT RNCs incubated in buffer for 16h and quantification of species. Representative gel of 3 biological repeats.

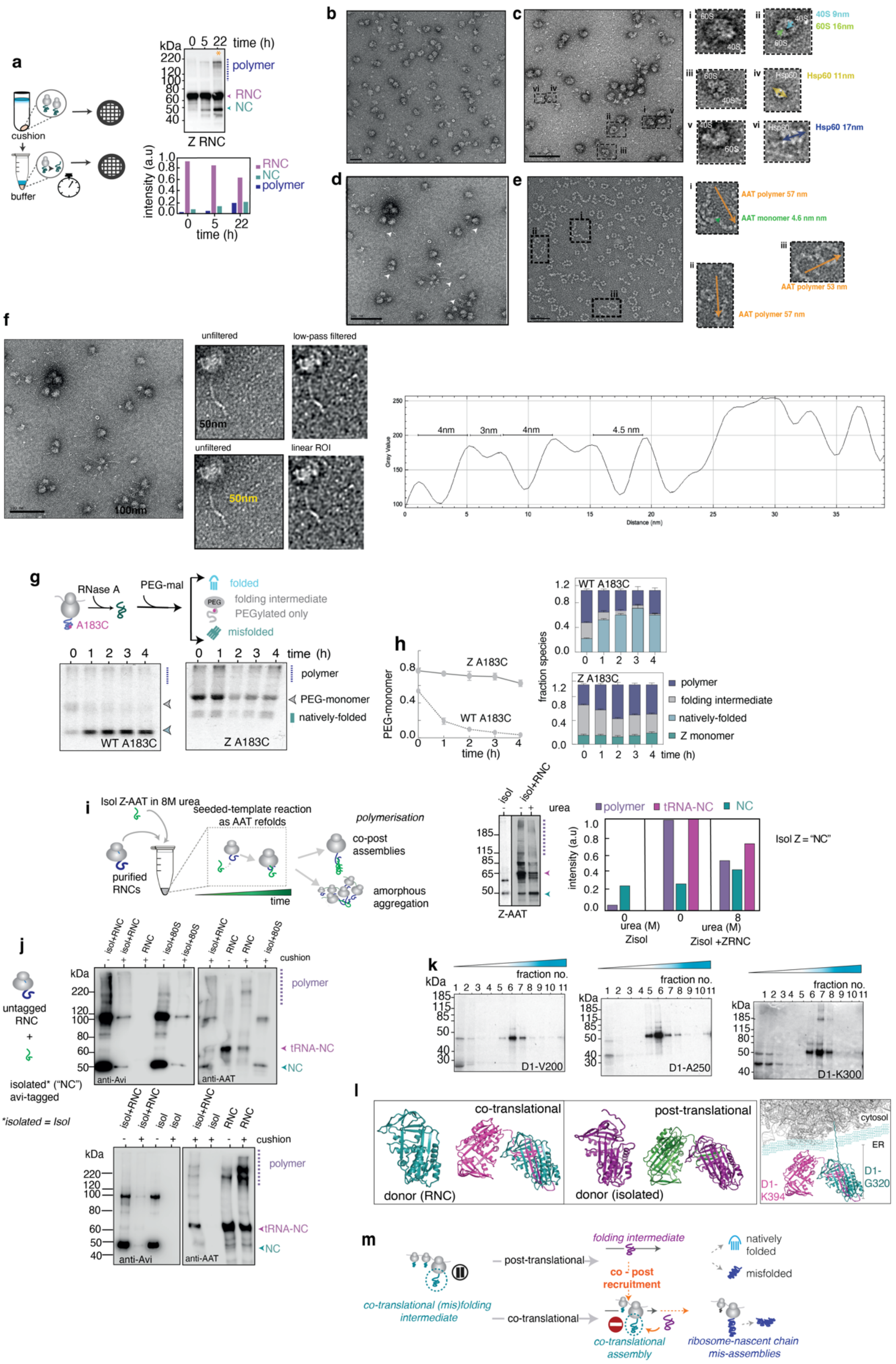


**Extended Data Figure 6: Characterisation of a co-post assembly mechanism (a)** *(left)* sample preparation used for negative-stain EM. *(right)* Time course of (a non-stop) Z-AAT RNC incubated over time. Shows stochastic release of AAT NCs and the accumulation of ribosome-bound and non-ribosome bound (released) polymers. **(b)** negative-stain micrograph of empty 80S ribosomes. **(c)** negative-stain micrograph of non-stop AAT RNCs produced in RRL and purified via a sucrose cushion. *(inset)* identifying species observed within the micrograph. **(d)** Full sized negative-stain micrograph of non-stop AAT RNCs as displayed in Fig. 3b. **(e)** negative-stain micrograph of purified (short) AAT polymers. **(f)** Analysis of AAT polymer lengths and AAT monomer units using Image J. **(g)** *(upper)* A PEGylation assay to measure the folding of A183C AAT NC following ribosome release with RNaseA treatment. *(lower)* Non-denaturing PAGE ($^{35}$S Met) showing the folding of released wild-type and Z A183C AAT NCs (n=3). **(h)** *(left)* The turnover of AAT's folding intermediate on-pathway to folded or misfolded states (n=3)**.** *(right)* Species formed during the folding of released AAT NCs (following RNaseA treatment) via PEGylation at A183C. **(i)** *(left)* A scheme showing how seeding is used to recapitulate a co-post assembly process *in vitro*. *(right)* Observation of co-post assembly following a seeding reaction in RRL and labelled with $^{35}$S methionine. **(j)** As in (i), but using unlabelled Z RNCs and seeding with Avi-tagged isolated Z AAT (anti-Avi and anti-AAT (1C2) western blot shown). **(k)** Co-translational misfolding and ribosome-bound polymer formation in AAT C-terminal truncations (D1-V200, D1-A250 and D1-K300). **(l)** Shows theoretical models of how dimers could form co- and post-translationally and in the ER. **(m)** A schematic model for a co-post mechanism of polymerisation.

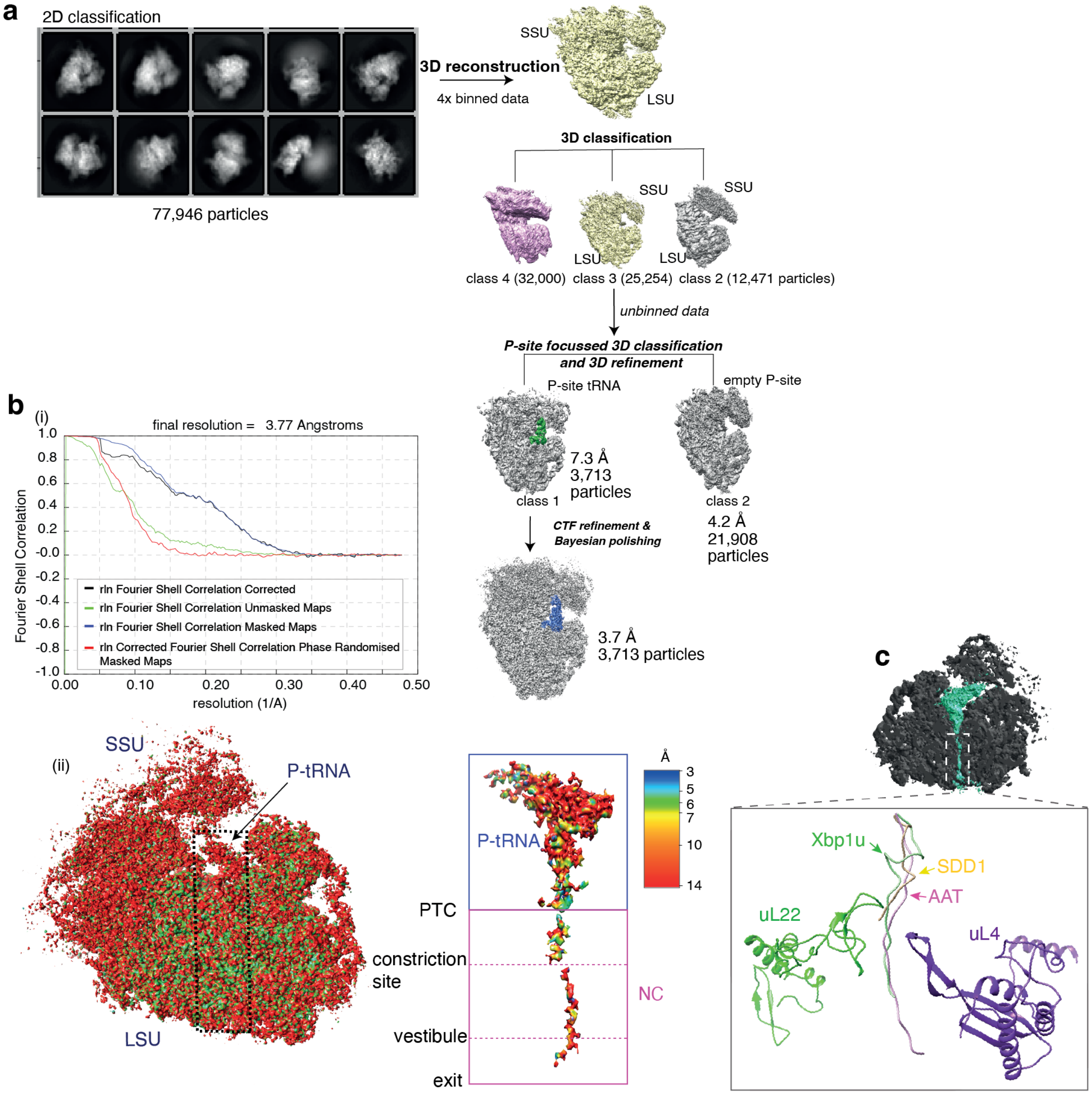


**Extended Data Figure 7: High resolution cryo-electron microscopy structure of wild-type AAT RNCs (a)** *(left)* Examples of 2D classes yielded after 2D classification of non-stop AAT RNC in Relion 3.1. *(right)* 3D classification of non-stop wild-type AAT RNCs performed in Relion 3.1. **(b)** *(i)* Fourier shell correlation curves extracted from post-processing in Relion 3.1. (ii) A 3D reconstruction of AAT RNCs with local resolution mapped to the ribosome and the emerging nascent chain as present within the ribosomal exit tunnel. **(c)** A superposition of the C-alpha model of AAT and two arrest-motifs, Xbp1u[10] and SDD1[65]. SSU = small subunit (40S), LSU = large subunit (60S)

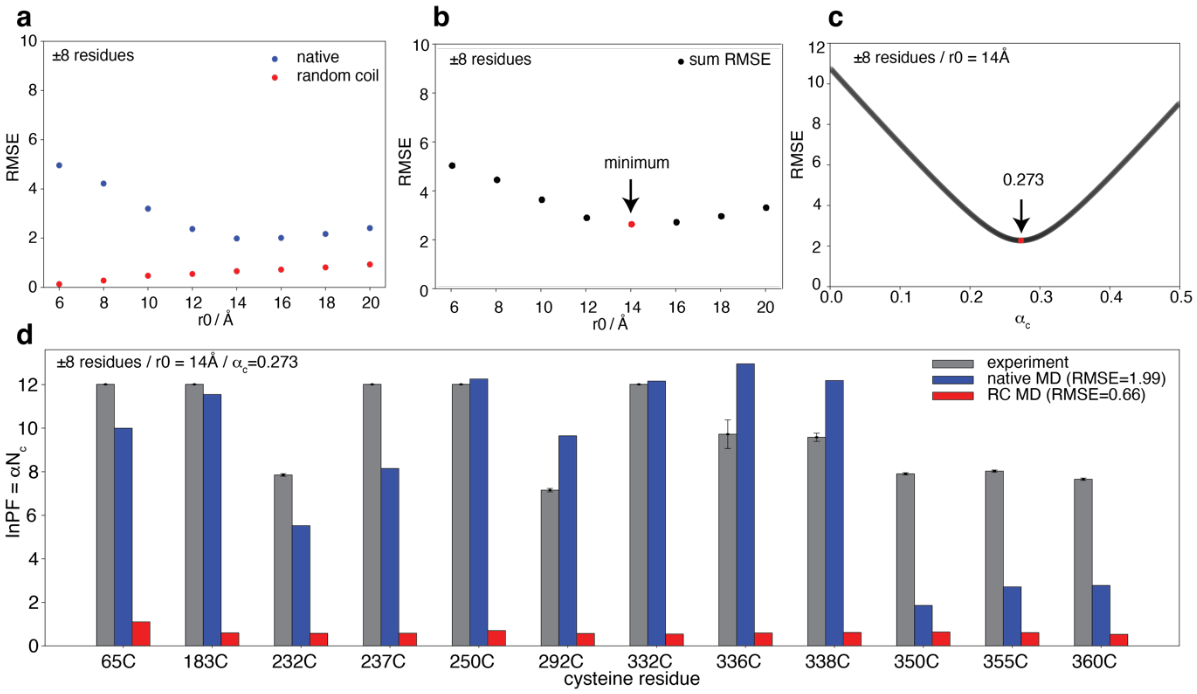


**Extended Data Figure 8: Parameter fitting for protection factor forward model (a)** Root Mean Square Error (RMSE) across series of r0 evaluating the fit of the native state ensemble to the experimental protection factors (blue) and the random coil ensemble fit to $\ln PF^{exp}=0$ (red) **(b)** Sum RMSE of native and random coil protection factor fits to the r0 at the minimum **(c)** RMSE of native state ensemble to experimental protection factors for a series of weights, **(d)** bar plot comparing the experimentally derived protection factors (grey) to the forward model ensemble predicted, and protection factors calculated for a fully disordered (random-coil, RC) model (blue).

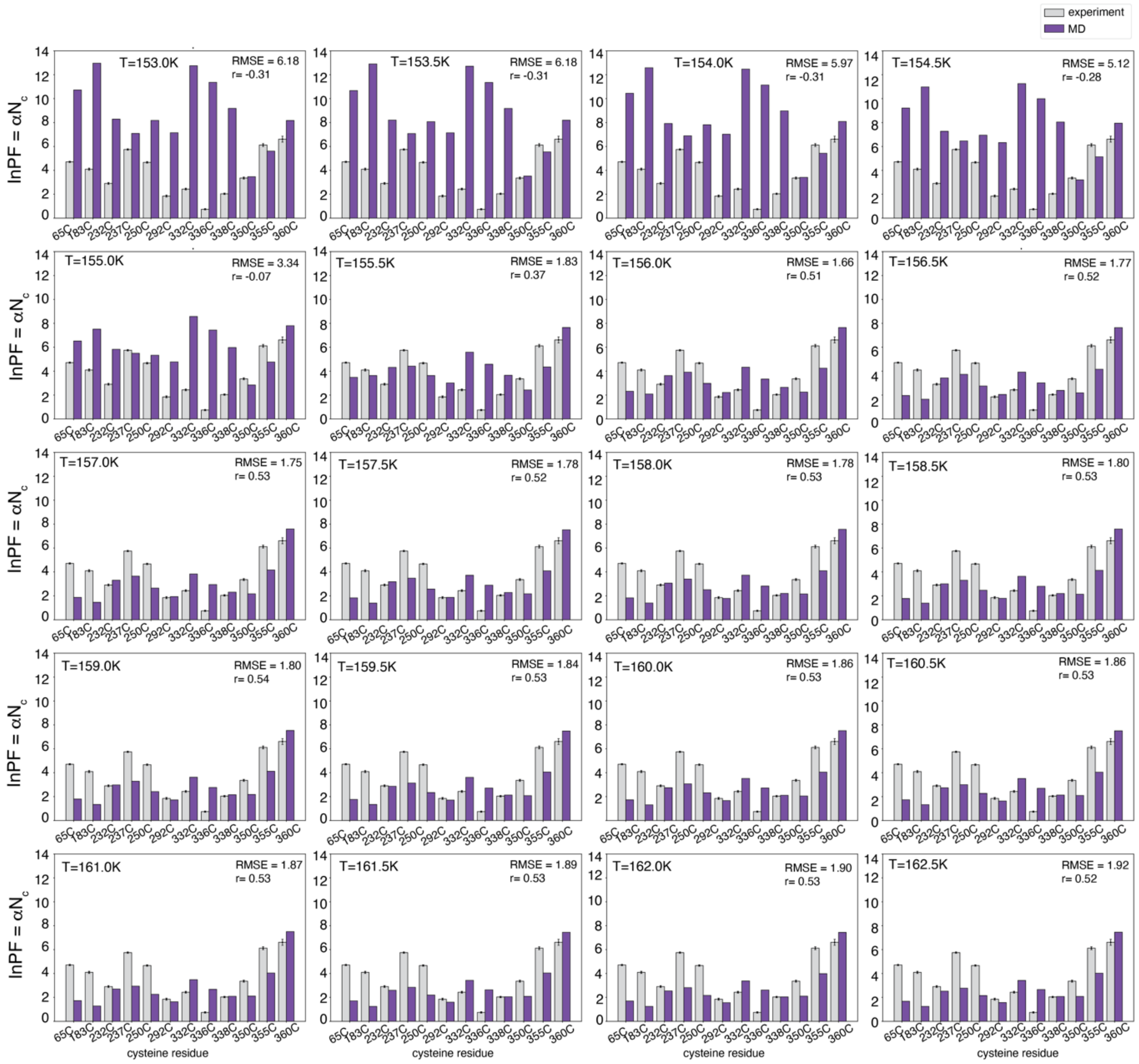

**Extended Data Figure 9: Prediction of protection factors from AAT RNC C-alpha structure-based model ensembles generated by PTMetaD**. Each window is a bar plot of protection factors predicted from the corresponding temperature replica (~45000 frames per replica). The ensemble at 156K indicates the ensemble with the lowest RMSE to experimental data and was the ensemble used for reweighting.

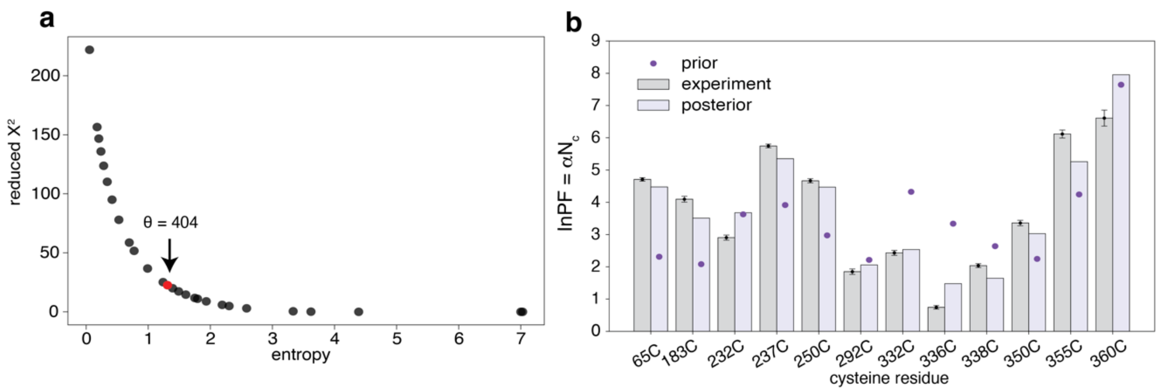


**Extended Data Figure 10: BioEn reweighting of AAT RNC ensemble with protection factor data (a)** L-curve analysis to choose confidence value **(b)** Plot of reweighted ensemble predicted protection factors, purple dots indicate the ensemble predicted protection factors prior to reweighting.

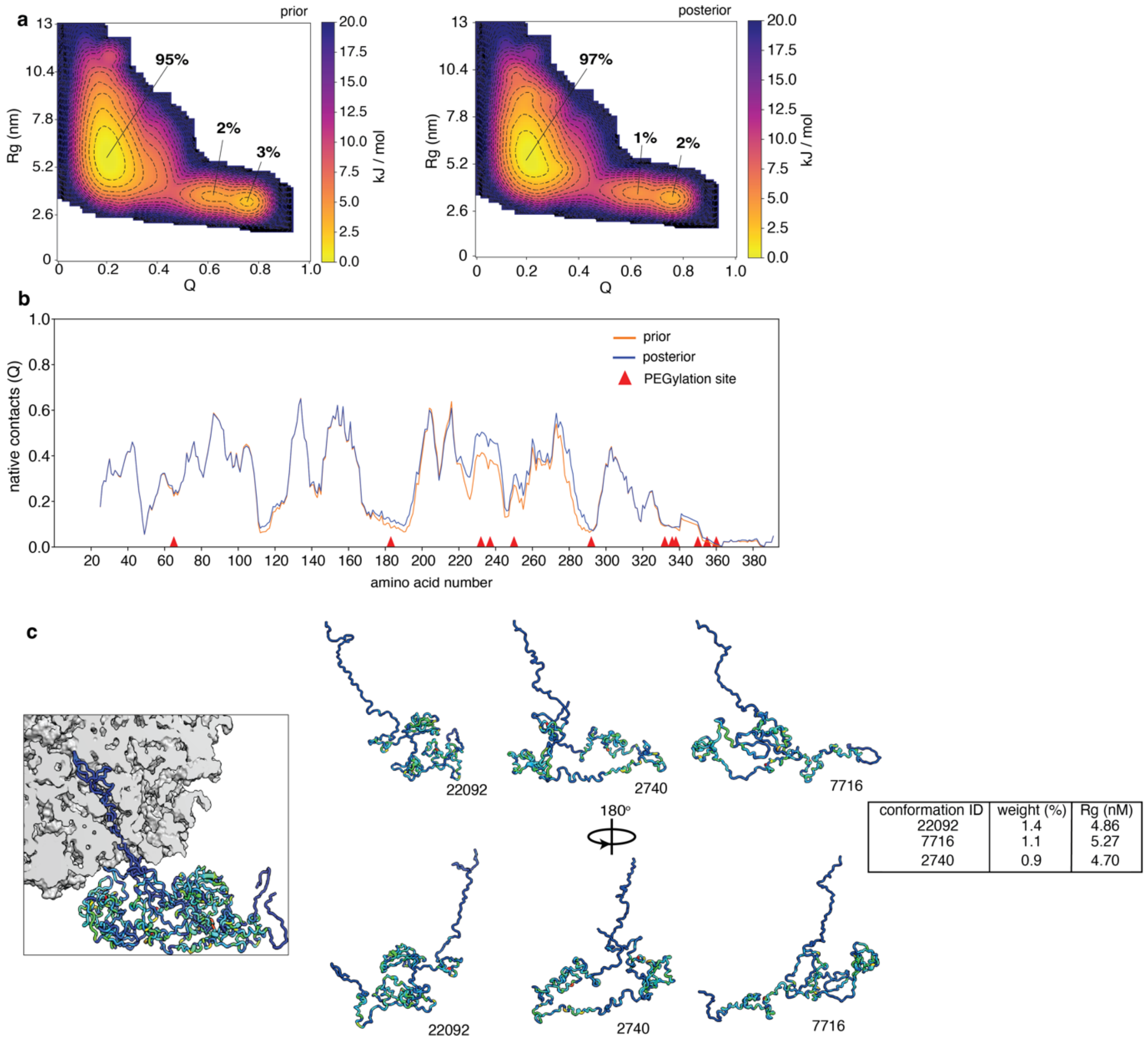


| conformation ID | weight (%) | Rg (nM) |
|---|---|---|
| 22092 | 1.4 | 4.86 |
| 7716 | 1.1 | 5.27 |
| 2740 | 0.9 | 4.70 |

**Extended Data Figure 11: Wild-type AAT co-translational folding intermediate ensemble (a)** A two-dimensional (2D) energy landscape depicting the distribution of the radius of gyration (Rg) values versus fraction of native contacts (Q), as observed for the conformational ensemble. Shown are energy landscapes before (prior) and after (posterior) reweighting. The majority of the wild-type AAT is in an expanded form, and two populations of highly-structured (native-like) intermediate (I-) states are also detected. These I-states show a significant native-state structure bias under these conditions used and were thus not studied further. The presence of structured I-states suggests that the steric effect of being ribosome-bound can still theoretically support native-like structure formation in AAT NCs. These highly structured I-states were observed both with and without experimental reweighting and likely contribute to I-state-like behaviour in the expanded species observed. **(b)** A weighted ensemble average fraction of native contacts (Q) per residue observed in the wild-type AAT co-translational folding intermediate before and after reweighting. **(c)** Overlay of three AAT NC structures (as shown in Fig. 3c(ii). *(right)* Individual conformations derived from Fig.3c(ii) (and from left), and a table of characteristics. Note that coarse-graining has been applied to both the 80S ribosome and the AAT NC. The NC occupying the tunnel shows apparent clashes with the 80S ribosome, however this is due to the choice of cartoon depiction only.

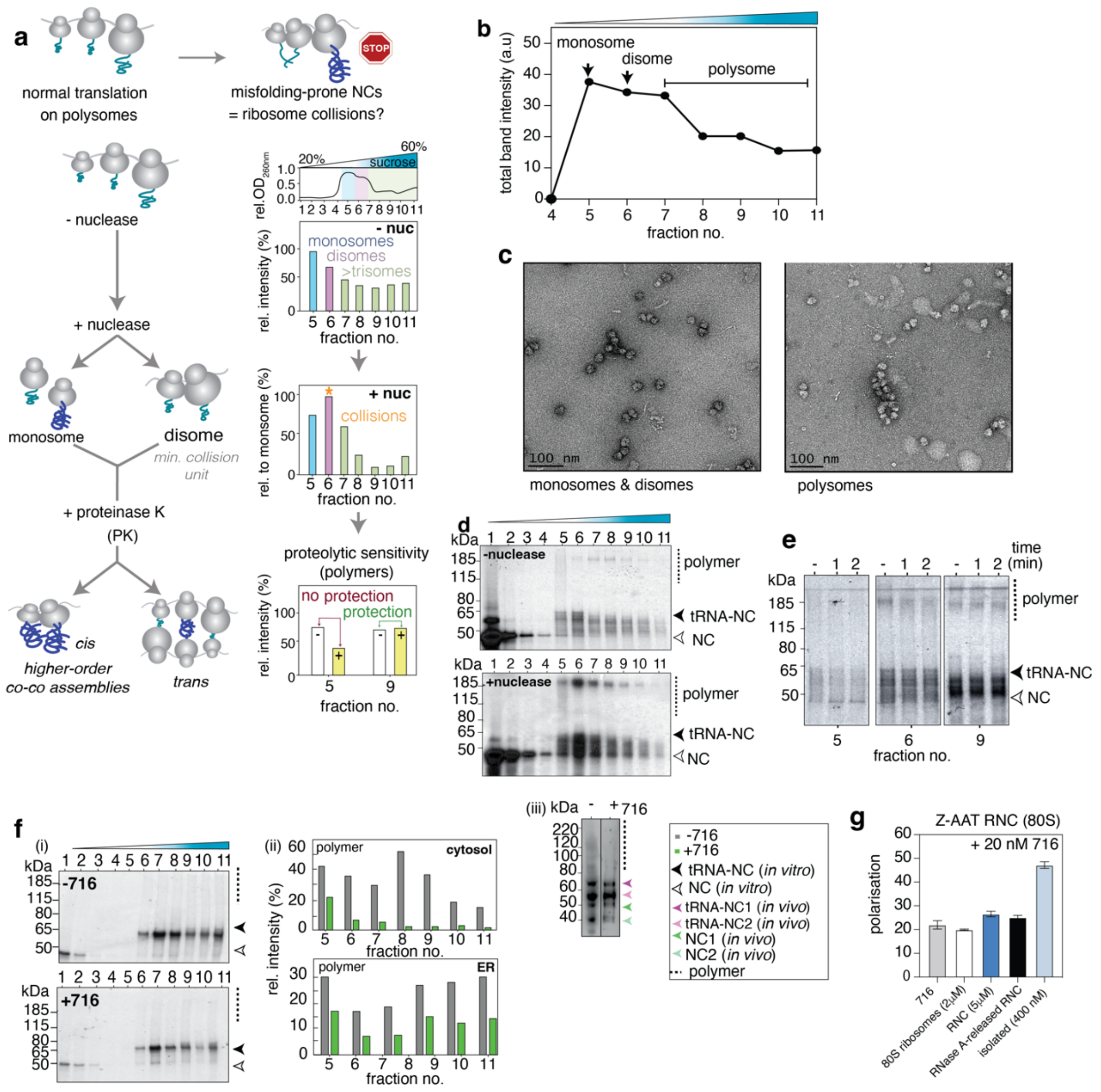

**Extended Data Figure 12: Characterisation of ribosome collisions via co-post assembly (a)** A schematic showing the experimental set-up to test for ribosome collisions and ribosome-bound assemblies. **(b)** Shows a sucrose gradient profile of Z-AAT determined using densitometry analysis (e.g., see (d, -nuclease)). **(c)** negative-stain micrographs of polysome sucrose fractions 7-11 showing monosomes, disomes and polysomes. **(d)** A sucrose gradient elution profile of Z-AAT (20 minutes biosynthesis), with and without nuclease treatment (n=4). **(e)** Proteinase K treatment of fractions 5 (monosomes), 6 (disomes) and 9 (polysomes) over time (n=3). **(f)** (i) The effect of 716 on polymer formation in ER microsomes (RRL). (ii) *(upper)* Quantification of the population of polymers in the absence and presence of 716 in the cytosol of RRL. *(lower)* Quantification of the population of polymers observed in ER microsomes in the absence and presence of 716. (iii) *(left)* The effect of 716 on polymer formation in HEK293 cells as observed in the ER (Xbpu1-stalled Z RNCs, 24 hours) (1C2 detection). **(g)** Fluorescence polarisation measurements of 80S ribosomes, RNCs, and isolated Z AAT samples incubated with 20 nM 716. Note that the average RNC occupancy in RRL is 30% thus the 5μM RNC sample has an effective NC concentration of 1.5μM.

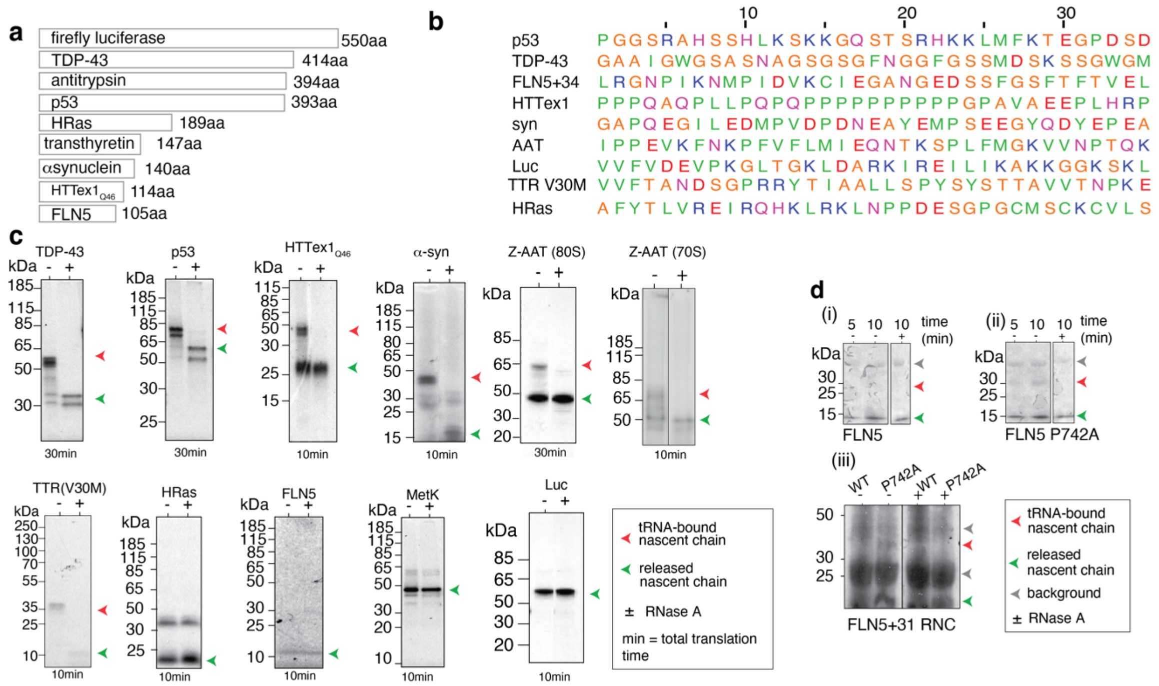


**Extended Data Figure 13: An energetic basis to persistent translational pausing (a)** Candidate proteins that were tested with the stalling assay. **(b)** Sequence alignment of the C-terminal 35 amino acids in the candidate proteins. **(c)** Extent of pausing observed across different systems as measured using $^{35}S$ methionine autoradiography (with longest time window shown for each). **(d)** (i-iii) Persistent translational pausing as observed in FLN5 and FLN5 P742A.
